# Protein *S*-acylation dynamics provide metabolic plasticity to acute myeloid leukemia cells

**DOI:** 10.64898/2026.03.02.708949

**Authors:** Nithya Balasundaram, Ayşegül Erdem, Azeem Sharda, Veerle W. Daniels, Phillip L. Chea, Fleur Leguay, Youzhong Liu, Mark A. Keibler, Charles Vidoudez, Andrew A. Lane, Didier Vertommen, Hans Casteur, Michaël R. Laurent, Sunia A. Trauger, Gregory Stephanopoulos, David T. Scadden, Nick van Gastel

## Abstract

Though cancer cells’ altered metabolism has been recognized for a century, the clinical success of metabolic targeting remains limited due to metabolic plasticity. Here, we use acute myeloid leukemia (AML) as a model to investigate this adaptability through combinatorial metabolic compound screening. Synthetic lethality emerged when AML cells were simultaneously treated with a glutaminase inhibitor and TOFA, a hypolipidemic agent. Sensitivity to this combination was also seen in primary patient samples and in other cancer types, while healthy hematopoietic progenitors were not affected. Unexpectedly, we discovered that TOFA acts through a non-canonical inhibition of protein *S*-acyltransferases. Protein *S*-acylation in AML cells specifically requires 16-to-18 carbon long fatty acids and is essential to maintain mitochondrial respiration upon glutaminolysis inhibition. Healthy cells in contrast have high intrinsic metabolic flexibility independent of *S*-acylation. Our results expose a unique mechanism of metabolic plasticity in cancer that could be targeted to enhance metabolic anti-cancer therapies.

## Introduction

Acute myeloid leukemia (AML) is an aggressive cancer of blood cells of the myeloid lineage, characterized by the clonal expansion of immature myeloid progenitors in the bone marrow and other hematopoietic tissues. While incidence of this disease increases with age, the response to treatment decreases, and 5-year survival is currently only 35%^1^. Despite recent advances in targeted therapies, AML remains a challenging cancer to treat, with relapse and treatment resistance being prominent obstacles to successful outcomes^2,3^. A growing body of evidence suggests that metabolic reprogramming plays a central role in the pathophysiology of AML, contributing to leukemogenesis, disease progression, and therapeutic resistance^4–7^. One key feature of AML metabolism is an increased dependency on mitochondrial oxidative phosphorylation (OXPHOS), observed in both AML blasts^8^ and leukemic stem cells (LSCs)^8,9^. While full inhibition of OXPHOS may be too toxic, as shown in Phase 1 clinical trials with the complex I inhibitor IACS-010759^10^, inhibition of individual metabolic pathways that fuel OXPHOS, such as glutaminolysis and fatty acid oxidation, can reduce AML progression, but not prevent it^11–15^.

Metabolic adaptability has emerged as a critical feature of cancer cell survival^16^. This includes both metabolic plasticity—the ability to process metabolic substrates in different ways—and metabolic flexibility—the ability to dynamically switch between different nutrients. In AML, the dysregulation of metabolic pathways is well-documented, but the mechanisms that enable leukemic cells to transition between these pathways and adapt in response to environmental changes and cellular demands are less clear. Moreover, emerging evidence suggests that while at diagnosis LSCs may have lower metabolic flexibility compared to AML blasts, the cells persisting through therapy and contributing to relapse acquire the ability to overcome metabolic interference^13^.

In the current study, we set out to investigate metabolic plasticity in AML using a combinatorial metabolic compound screening approach. We discovered an unexpected synthetic lethality between bis-2-(5-phenylacetamido-1,2,4-thiadiazol-2-yl)ethyl sulfide (BPTES), an inhibitor of glutaminolysis, and 5-(tetradecyloxy)-2-furoic acid (TOFA), a fatty acid analog and hypolipidemic agent. Detailed mechanistic studies led to the identification of protein *S*-acylation as a non-canonical TOFA target and crucial mechanism allowing AML cells to overcome the inhibition of glutaminolysis. Importantly, this dependency is not shared by healthy hematopoietic progenitor cells, which possess a much larger inherent metabolic flexibility, but may be used also by other cancer types, highlighting the therapeutic potential of our metabolic inhibitor combination.

## Results

### Combined treatment with BPTES and TOFA selectively kills AML cells

Based on the observation that AML cells are generally able to overcome the blockade of a single metabolic pathway, we hypothesized that combined inhibition of two metabolic pathways could result in unexpected synthetic lethality. To test this, we set up an *in vitro* combinatorial screen with 11 metabolic compounds targeting enzymes across central carbon metabolism, involved in the pentose phosphate pathway (6-aminonicotinamide, 6AN), glycolysis (TEPP46; FX11), glucose oxidation (dichloroacetate, DCA; UK5099), oxidative phosphorylation (phenformin, PHEN), glutaminolysis (BPTES; epigallocatechin gallate, EGCG), fatty acid oxidation (etomoxir), *de novo* lipogenesis (TOFA) and transamination (aminooxyacetate, AOA) (Figure S1A). We first performed dose-response experiments using healthy human bone marrow-derived mononuclear cells (BM-MNCs) and cord blood-derived CD34^+^ cells (CB-CD34^+^ cells) to determine the highest concentration of each compound that does not induce cell death in normal hematopoietic stem and progenitor cells (HSPCs) (Figure S1B). Next, we exposed four different human AML cell lines (NOMO1, NB4, U937, MOLM14) to all possible combinations of two compounds at the selected concentrations and evaluated viability after 72 hours. While individual cell lines showed specific sensitivities, the combination of BPTES and TOFA showed efficient and synergistic cell death induction across all four cell lines (Figure 1A).

**Figure 1.**
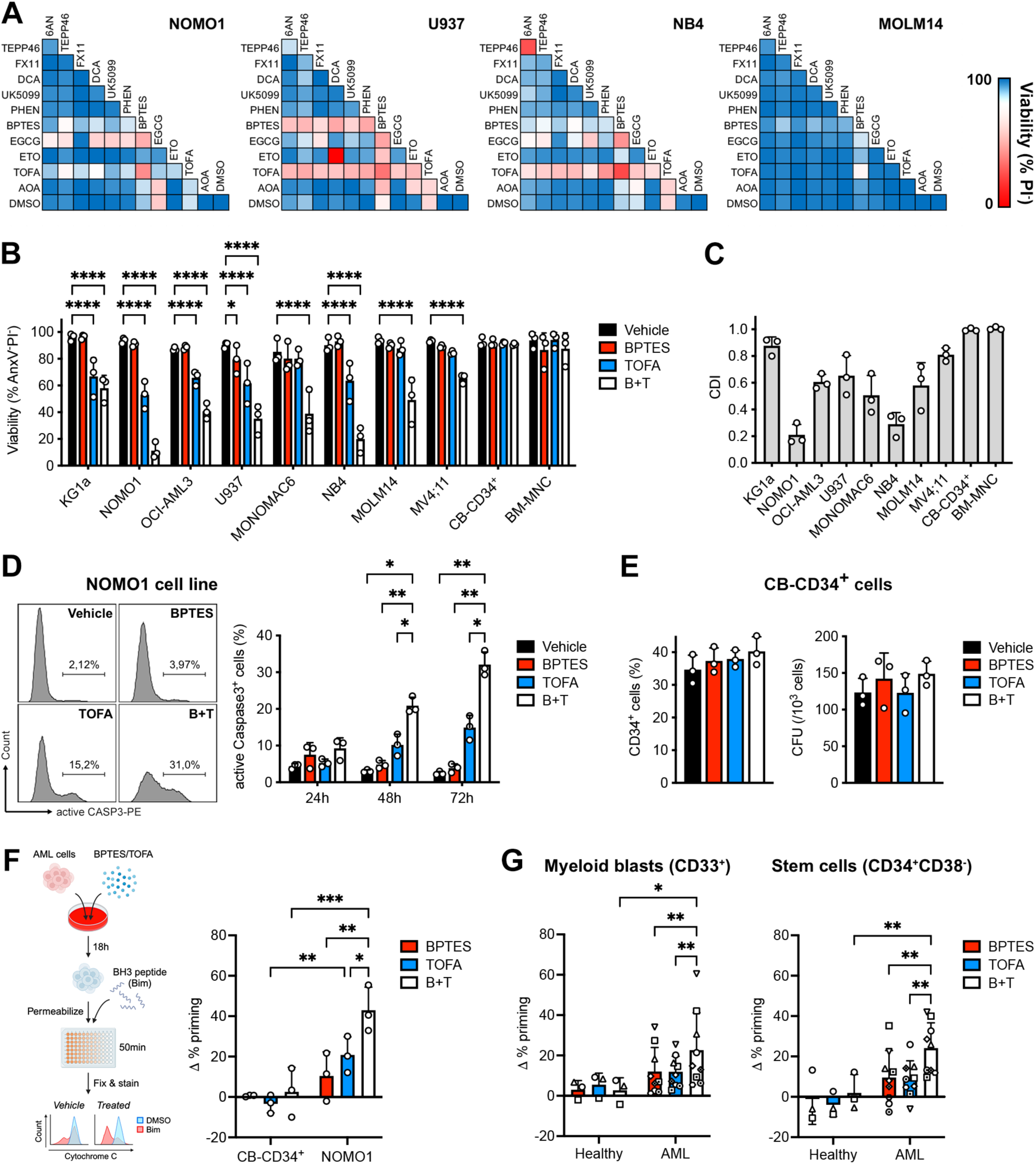
Synthetic Lethality of BPTES and TOFA in AML. (**A**) Heatmaps showing viability of AML cell lines treated for 72 hours with combinations of different metabolic inhibitors (see Figure S1B for inhibitor concentrations). PI, propidium iodide. (**B**) Viability of AML cell lines and healthy hematopoietic stem/progenitor cells treated for 72 hours with BPTES (5 µM) and/or TOFA (5 µM). CB-CD34^+^, cord blood-derived CD34^+^ cells; BM-MNC, bone marrow mononuclear cells. (**C**) Coefficient of Drug Interaction (CDI) for BPTES and TOFA calculated from the viability data shown in B. (**D**) Percentage of apoptotic (active Caspase 3^+^) NOMO1 AML cells at different time points after treatment with BPTES and/or TOFA. Plots on the left show a representative sample at 72 hours. (**E**) Percentage of CD34^+^ cells (left) and colony-forming capacity (right) of CB-CD34^+^ cell cultures treated for 72 hours with BPTES and/or TOFA. CFU, colony-forming unit. (**F-G**) Apoptotic priming of CB-CD34^+^ HSPCs and NOMO1 AML cells (F) or primary healthy (3 donors) and AML patient (9 donors) bone marrow myeloid blasts and stem cells (G) treated for 18 hours with BPTES and/or TOFA, as determined by BH3 profiling and expressed relative to vehicle controls. Data are represented as mean ± SD. *p < 0.05, **p < 0.01, ***p < 0.001, ****p < 0.0001. See also Figure S1.

We validated these findings using eight different human AML cell lines, all of which proved highly sensitive to combined BPTES and TOFA treatment, while healthy BM-MNCs or CB-CD34^+^ cells did not show any signs of cell death (Figure 1B, Figure S1C-D). Calculation of the coefficient of drug interaction (CDI) showed that induction of cell death by the combination of BPTES and TOFA is synergistic (CDI<1), with most cell lines showing highly synergistic effects (CDI<0.7) (Figure 1C, Figure S1E). This synthetic lethality was seen across compound concentrations (Figure S1F). Cell death in AML cells exposed to BPTES and TOFA occurred through apoptosis, as suggested by Annexin V staining (Figure 1B, Figure S1C) and confirmed by detection of active Caspase 3 as early as 48 hours after treatment (Figure 1D). To further validate that non-malignant HSPCs were not affected by the drug combination, we treated cells for 72 hours with BPTES and/or TOFA, and checked the percentage of primitive CD34^+^ cells and the ability of the cells to form colonies, both of which were unaffected (Figure 1E).

We next sought to confirm the cytotoxic effects of the combination of BPTES and TOFA on primary AML patient-derived cells. Given the difficulty in culturing primary AML patient samples *in vitro* for prolonged times^17^, we decided to perform dynamic BH3 profiling (DBP), a technique that can predict cytotoxicity by measuring early drug-induced changes in mitochondrial apoptotic priming and which requires the cells to be cultured for a short time only^18,19^. We first confirmed that DBP adequately predicts the sensitivity of the NOMO1 AML cell line to BPTES and TOFA, and the absence of apoptosis in CB-CD34^+^ cells exposed to the inhibitors (Figure 1F). Next, we collected bone marrow samples from AML patients and healthy controls (Table S1), isolated mononuclear cells and exposed them for 18 hours to BPTES and/or TOFA, after which DBP was performed. Like observed for the NOMO1 cell line, BPTES and TOFA increased mitochondrial apoptotic priming in both the CD33^+^ myeloid blast fraction and the stem cell-enriched CD34^+^CD38^-^ fraction of the AML patient cells, with the combination showing a significantly higher effect than either compound alone (Figure 1G). Together, these results show that the combination of BPTES and TOFA induces synthetic lethality in AML cells, but not in healthy HSPCs.

### AML cells show hyperactivation of *de novo* lipogenesis

We next set out to validate the targets of both metabolic inhibitors. BPTES is known as an inhibitor of glutaminase (GLS)^20^, while TOFA inhibits the activity of acetyl-CoA carboxylase (ACC)^21^. *GLS*, but not *GLS2*, was expressed in all AML cell lines as well as in CB-CD34^+^ cells, although protein levels for GLS were much higher in AML compared to non-malignant cells (Figure 2A, Figure S2A). For ACC, cells mainly expressed ACC1, the cytoplasmic isoform involved in *de novo* lipogenesis, while levels of ACC2, the mitochondrial isoform known to regulate fatty acid oxidation, were lower at both mRNA and protein levels. Again, protein analysis showed that while differences between individual cell lines exist, AML cell lines have overall much higher levels of ACC1 compared to CB-CD34^+^ cells (Figure 2A, Figure S2A).

**Figure 2.**
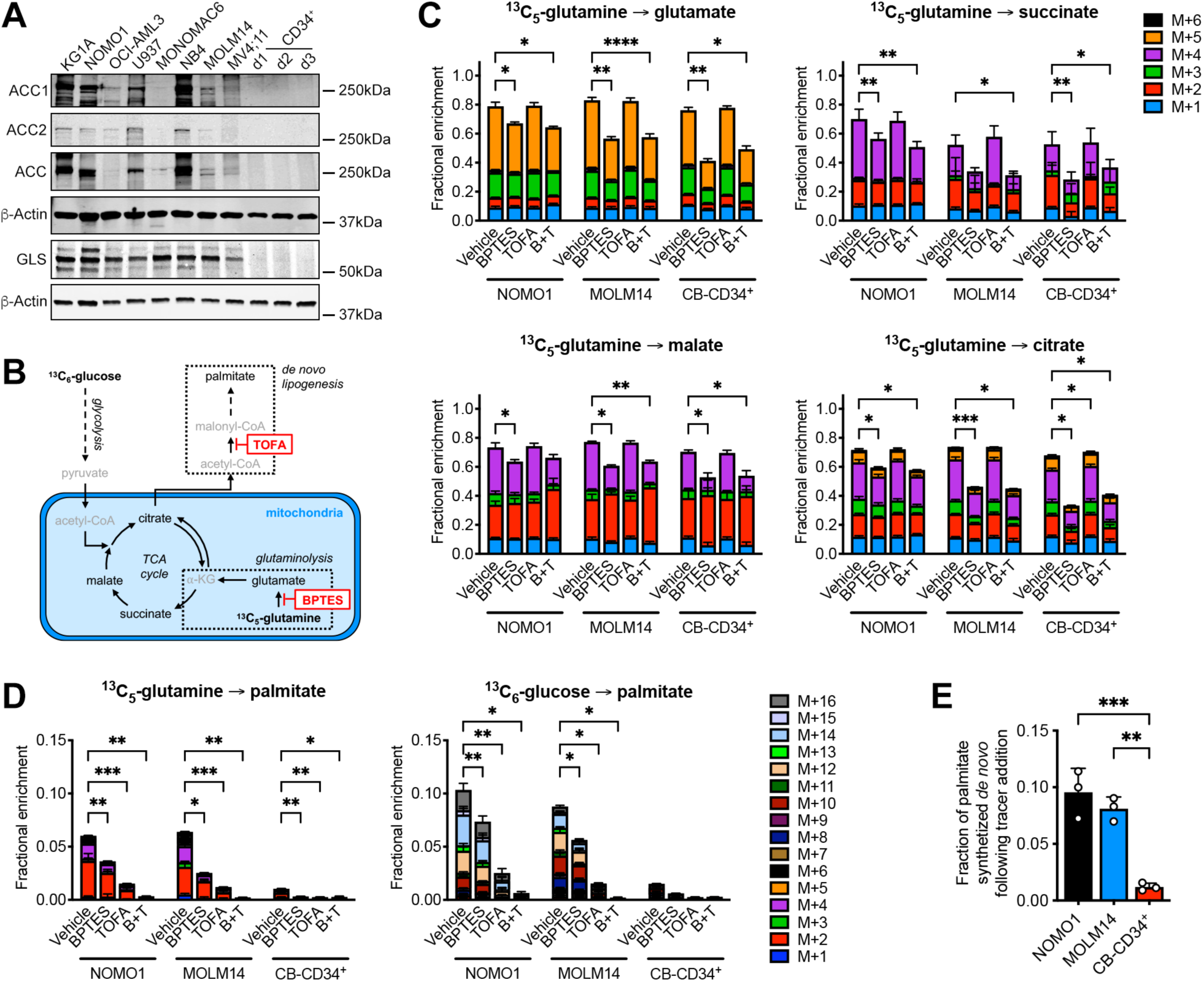
AML Cells Show Hyperactivation of *De Novo* Lipogenesis. (**A**) Immunoblot detection of acetyl-CoA carboxylase 1 (ACC1), ACC2, total ACC and glutaminase (GLS) protein levels in AML cell lines or cord blood-derived CD34^+^ cells (three independent donors shown), with β-actin as loading control. (**B**) Schematic overview of the stable isotope tracing experiments. (**C**) Fractional enrichment of ^13^C_5_-glutamine in intracellular glutamate, succinate, malate and citrate in AML cell lines or cord blood-derived CD34^+^ cells (CB-CD34^+^) treated for 24 hours with BPTES (5 µM) and/or TOFA (5 µM). Statistical analysis was performed by two-way ANOVA on the sum of all isotopomers. (**D**) Fractional enrichment of ^13^C_5_-glutamine or ^13^C_5_-glucose in intracellular lipid palmitate in AML cell lines or CB-CD34^+^ treated for 24 hours with BPTES and/or TOFA. Statistical analysis was performed by two-way ANOVA on the sum of all isotopomers. (**E**) Fraction of intracellular lipid palmitate synthetized from glucose in AML cell lines or CB-CD34^+^ cells over the course of 24 hours of treatment with BPTES and/or TOFA. Data are represented as mean ± SD. *p < 0.05, **p < 0.01, ***p < 0.001, ****p < 0.0001. See also Figure S2.

To confirm that BPTES and TOFA induced the expected metabolic changes, we performed stable isotope metabolic tracer analysis (Figure 2B). Treatment of NOMO1 or MOLM14 AML cells with BPTES reduced labeling of glutamate and TCA cycle metabolites from uniformly ^13^C-labeled glutamine, in line with inhibition of GLS (Figure 2C), and accordingly reduced cellular levels of glutamate, succinate, malate and citrate while increasing glutamine pools (Figure S2B). The labeling patterns for CB-CD34^+^ cells at baseline or after BPTES treatment were very similar to those of the AML cells (Figure 2C, Figure S2B), indicating comparable glutaminolysis activity despite very different levels of GLS (Figure 2A). TOFA did not have any effects on glutaminolysis in AML or CB-CD34^+^ cells, neither as a single agent nor in combination with BPTES (Figure 2C).

To assess *de novo* lipogenesis, we traced carbon from uniformly ^13^C-labeled glutamine or glucose to palmitate (Figure 2B). Analysis of labeling patterns at baseline showed that AML cells synthetize palmitate predominantly from glucose carbons (enrichment of M+10, M+12, M+14 and M+16 isotopomers), with only minor contribution from glutamine (mainly M+2 and M+4 isotopomers) (Figure 2D). In accordance, reductive carboxylation of glutamine, which often occurs when glutamine carbons are used for *de novo* lipogenesis, was low compared to oxidative glutamine catabolism, as shown by low levels of M+5 versus M+4 citrate (Figure 2C). BPTES slightly decreased labeling of palmitate, while TOFA strongly reduced palmitate synthesis from both glucose- and glutamine-derived carbons, as expected (Figure 2D). The combination of BPTES and TOFA decreased *de novo* lipogenesis more extensively compared to either compound alone. Despite the profound inhibition of *de novo* lipogenesis, we did not detect changes in total cellular palmitate pools upon treatment of AML cells with BPTES and/or TOFA (Figure S2B).

The rate of *de novo* lipogenesis was much lower in CB-CD34^+^ cells compared to AML cells, as evidenced by the almost complete lack of labeling in palmitate from either glucose or glutamine (Figure 2D). Using isotopomer spectral analysis we calculated the fraction of palmitate newly synthesized from glucose after tracer addition, which showed that CB-CD34^+^ cells have a rate of fatty acid synthesis that is 8-to-10-fold lower than that of AML cells (Figure 2E). Despite this low basal rate, treatment with BPTES and/or TOFA further decreased *de novo* lipogenesis in HSPCs to almost undetectable levels (Figure 2D).

These data thus show that both BPTES and TOFA have the expected metabolic effects in AML cells and HSPCs, and that AML cells exhibit hyperactivation of *de novo* lipogenesis compared to HSPCs.

### AML cells require long chain saturated and monounsaturated fatty acids

Since treatment with BPTES and TOFA led to a strong decrease in *de novo* lipogenesis, we investigated how this impacts AML cell lipid metabolism. We detected increased fatty acid uptake in different AML cell lines and HSPCs after treatment, with the highest increase induced by TOFA alone or in combination with BPTES (Figure 3A; Figure S3A), which may explain why we did not observe changes in total cellular palmitate pools after TOFA treatment (Figure S2B). However, the response in extracellular lipid uptake after BPTES and/or TOFA treatment differed between individual AML cell lines (Figure 3A) and did not correlate with the basal level of lipid uptake (Figure 3B) or the sensitivity to TOFA treatment (Figure 1B).

**Figure 3.**
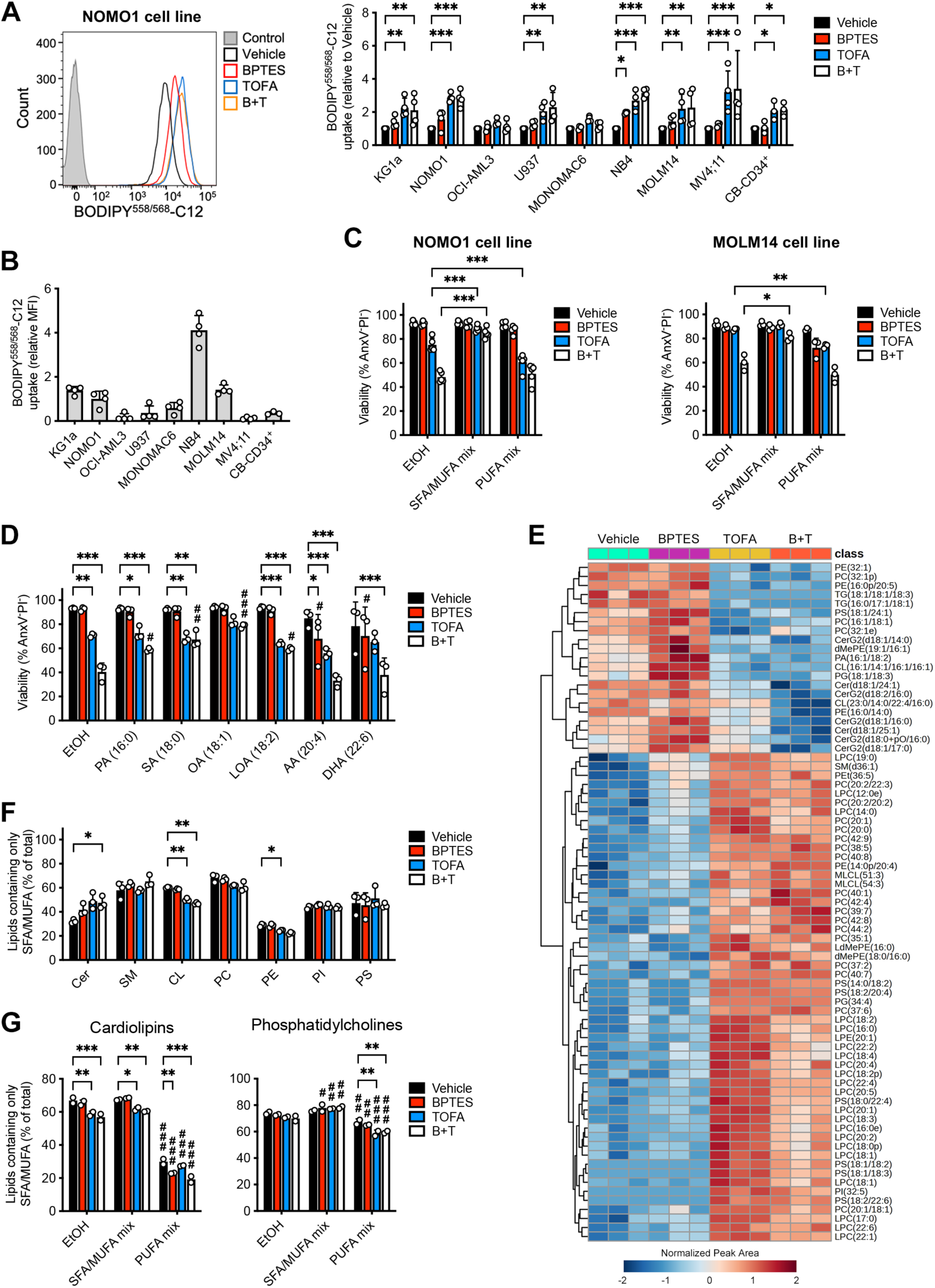
AML Cells Require Saturated and Monounsaturated Fatty Acids. (**A**) Uptake of a fluorescently labeled C12 fatty acid in AML cell lines or cord blood-derived CD34^+^ cells (CB-CD34^+^) treated for 24 hours with BPTES (5 µM) and/or TOFA (5 µM), relative to vehicle-treated cells. Plot on the left shows a representative sample of the NOMO1 cell line, with unlabeled cells included as Control. (**B**) Comparison of fluorescently labeled C12 fatty acid uptake across different AML cell lines and CB-CD34^+^ cells at baseline. (**C**) Viability of AML cell lines treated for 72 hours with BPTES and/or TOFA, in the presence of exogenously added mixtures of albumin-complexed saturated/monounsaturated fatty acids (SFA/MUFA; palmitic acid, stearic acid and oleic acid at 15 µM each) or polyunsaturated fatty acids (PUFA; linoleic, arachidonic and docosahexaenoic acid at 15 µM each). Ethanol/albumin (EtOH) was used as vehicle. (**D**) Viability of NOMO1 AML cells treated for 72 hours with BPTES and/or TOFA, in the presence of exogenously added albumin-complexed fatty acids (15 µM). PA, palmitic acid; SA, stearic acid; OA, oleic acid; LOA, linoleic acid; AA, arachidonic acid; DHA, docosahexaenoic acid. (**E**) Heatmap showing the 75 most-changed lipid species (ANOVA) in NOMO1 AML cells treated for 24 hours with BPTES and/or TOFA, as determined by lipidomics. (**F**) Saturation degree across different lipid classes in NOMO1 AML cells treated for 24 hours with BPTES and/or TOFA, as determined by lipidomics. Cer, ceramides; SM, sphingomyelins; CL, cardiolipins; PC, phosphatidylcholines; PE, phosphatidylethanolamines; PI, phosphatidylinositols; PS, phosphatidylserines. (**G**) Saturation degree of CL and PC phospholipids in NOMO1 AML cells treated for 24 hours with BPTES and/or TOFA, in the presence or absence of exogenously added SFA/MUFA or PUFA mixtures. Data are represented as mean ± SD. *p < 0.05, **p < 0.01, ***p < 0.001 versus Vehicle; ^#^p < 0.05, ^##^p < 0.01, ^###^p < 0.001 versus EtOH. See also Figure S3.

We then tested whether supplementation with additional exogenous fatty acids could rescue the effects of BPTES and TOFA treatment on AML cell viability. Adding a mixture of saturated and monounsaturated fatty acids (SFAs/MUFAs; palmitic acid, stearic acid, and oleic acid), representing the fatty acids that can be produced through *de novo* lipogenesis, led to an almost complete rescue of cell death induction (Figure 3C). In contrast, addition of polyunsaturated fatty acids (PUFAs; linoleic acid, arachidonic acid, and docosahexaenoic acid), which cannot be produced by mammalian cells and need to be taken up from the environment, did not affect viability. When testing exogenous supplementation of individual fatty acids, we found that 16-to-18 carbon long fatty acids, and particularly oleic acid, were the most effective at preventing TOFA-induced cell death (Figure 3D). Supplementation with very long chain PUFAs such as arachidonic acid or docosahexaenoic acid in contrast slightly increased cell death when combined with TOFA (Figure 3D).

To better characterize the effects of BPTES and TOFA on the cellular lipid composition, we next performed untargeted lipidomics analysis on the NOMO1 AML cells. We detected 1700-1800 total lipid species per independent experiment, of which 1088 unique species, from 31 lipid classes, were consistently detected across experiments and used for further analysis (Table S2). When comparing the overall lipidome by Principal Component Analysis (Figure S3B), we observed a large shift induced by TOFA, while BPTES had only minor effects. TOFA led to an overall depletion of triglycerides and increase in lysophospholipids (Figure 3E, Figure S3C). This indicates that besides increasing fatty acid uptake, AML cells also mobilize intracellular lipids stored as triglycerides when *de novo* lipogenesis is inhibited. However, the observed increase in lysophospholipids, in addition to the exogenous fatty acid supplementation experiments described above, confirms that these adaptations are not sufficient to maintain the overall SFA/MUFA pools in TOFA-treated cells. In accordance, lipids containing at least one SFA/MUFA tail (myristic acid – C14:0, myristoleic acid – C14:1, palmitic acid – C16:0, palmitoleic acid – C16:1, stearic acid – C18:0, or oleic acid – C18:1) were particularly depleted in AML cells treated with TOFA alone or in combination with BPTES, while lipids containing very long PUFA tails were highly increased (Figure 3E). When examining the overall saturation degree across different lipid classes, calculated as the fraction of lipid species in each class that is composed exclusively of SFAs and MUFAs (Figure 3F, Figure S3D), we found the most striking decrease in saturation degree after TOFA treatment in cardiolipins, with smaller effects in phosphatidylcholine, phosphatidylethanolamine, phosphatidylserine, phosphatidylglycerol and phosphatidic acid species. Ceramides and monolysocardiolipins in contrast showed an increase in saturation degree after TOFA.

Our results show that inhibition of *de novo* lipogenesis by TOFA deprives AML cells of SFAs and MUFAs, promoting membrane lipid unsaturation by increased incorporation of PUFAs. Prior studies in melanoma, prostate, lung, and breast cancer have shown that increased membrane lipid unsaturation can sensitize cancer cells to lipid peroxidation and promote oxidative stress-induced cell death and ferroptosis^22–25^. Given that inhibition of GLS in AML cells is known to increase cellular reactive oxygen species (ROS) levels^26^, we investigated whether the cell death induced by the combination of BPTES and TOFA is linked to oxidative stress and/or lipid peroxidation. While we could confirm that inhibition of GLS by BPTES increases total cellular ROS levels as well as mitochondrial superoxide levels in AML cells (Figure S3E-F), we found no evidence of increased lipid peroxidation in cells treated with the combination of BPTES and TOFA (Figure S3G). In line with this, the ferroptosis inhibitor Ferrostatin-1 had no effect on AML cell death induced by BPTES and TOFA, although it efficiently prevented cell death induced by the known ferroptosis-inducing agent Erastin (Figure S3H). Together with the detection of active Caspase 3 (Figure 1D) and increased apoptotic priming (Figure 1F-G), these results confirm that AML cells treated with TOFA alone or in combination with BPTES undergo apoptotic and not ferroptotic cell death.

We next focused on cardiolipins, a type of phospholipid containing four acyl chains that is almost exclusively found in mitochondria, where it plays key roles in both energy metabolism and apoptosis^27,28^. A key step in apoptosis induction is cardiolipin oxidation, a process that can only occur at unsaturations in the fatty acid tails. As TOFA induces cardiolipin unsaturation (Figure 3F), and BPTES increases mitochondrial ROS levels (Figure S3F), we questioned whether cardiolipin oxidation lays at the basis of the synergistic cell death induction between the two compounds. This was however not the case, as cardiolipin-specific 10-N-nonyl acridine orange (NAO) staining, which is known to decrease upon cardiolipin oxidation^29^, showed no change with TOFA and even a small increase in cells treated with BPTES alone (Figure S3I), perhaps reflecting a subtle increase in overall cardiolipin pools in BPTES-treated cells. In addition, we detected several oxidized and peroxidized cardiolipin species in our lipidomics dataset, but the levels of these were not increased in cells treated with TOFA or the combination of BPTES and TOFA (Figure S3J).

To better understand whether the observed AML cell apoptosis is in any way related to the increased membrane unsaturation induced by TOFA, we performed lipidomics analysis on cells treated with BPTES and/or TOFA in the presence or absence of the mixtures of SFAs/MUFAs or PUFAs (Table S3), as these fatty acid mixes differed in their ability to rescue BPTES/TOFA-induced AML cell death (Figure 3C). Supplementation of cells with either the SFA/MUFA or PUFA mix restored total triglyceride levels and lysophospholipid levels of TOFA-treated cells nearly back to normal, showing that these treatments can largely prevent cellular fatty acid depletion (Figure S3K). A strong increase in total acylcarnitine levels was also observed in cells supplemented with exogenous fatty acids, indicating an increase in fatty acid oxidation. However, the effects of the SFA/MUFA or PUFA mix on total lipid levels across classes were nearly identical. Analysis of the saturation degree showed that supplementation with SFAs/MUFAs slightly increased the saturation degree in most lipid classes (including cardiolipins, phosphatidylcholines, phosphatidylserines, and triglycerides) in cells treated with TOFA, although this was not fully normalized to the level of vehicle-treated cells (Figure 3G, Figure S3L). In contrast, supplementation with exogenous PUFAs led to a massive decrease in lipid saturation in many classes, including cardiolipins, phosphatidylcholines, phosphatidylethanolamines, phosphatidylinositols, phosphatidylserines, phosphatidylglycerols, lysophospholipids, triglycerides and acylcarnitines, irrespective of whether cells were treated with metabolic inhibitors or not (Figure 3G, Figure S3K). Yet, despite this massive membrane lipid polyunsaturation, supplementation of AML cells with the PUFA mixture did not induce apoptosis or sensitization to BPTES (Figure 3C).

Taken together, these results show that while the effects of TOFA on AML cell viability are related to a process requiring long chain SFAs and MUFAs, they are independent of its effects on membrane lipid saturation.

### The pro-apoptotic effects of TOFA are not due to its inhibition of *de novo* lipogenesis

To better understand the mechanism by which BPTES and TOFA synergize, we next tested whether the cytotoxic effects of these molecules could be fully ascribed to their respective inhibition of GLS and ACC. When BPTES was replaced by either Compound 968 (C968) or CB-839, two unrelated GLS inhibitors, a similar synergy was observed with TOFA in reducing viability of the NOMO1 and MOLM14 cell lines (Figure 4A-B), thus confirming that the effects of BPTES are due to its inhibition of GLS.

**Figure 4.**
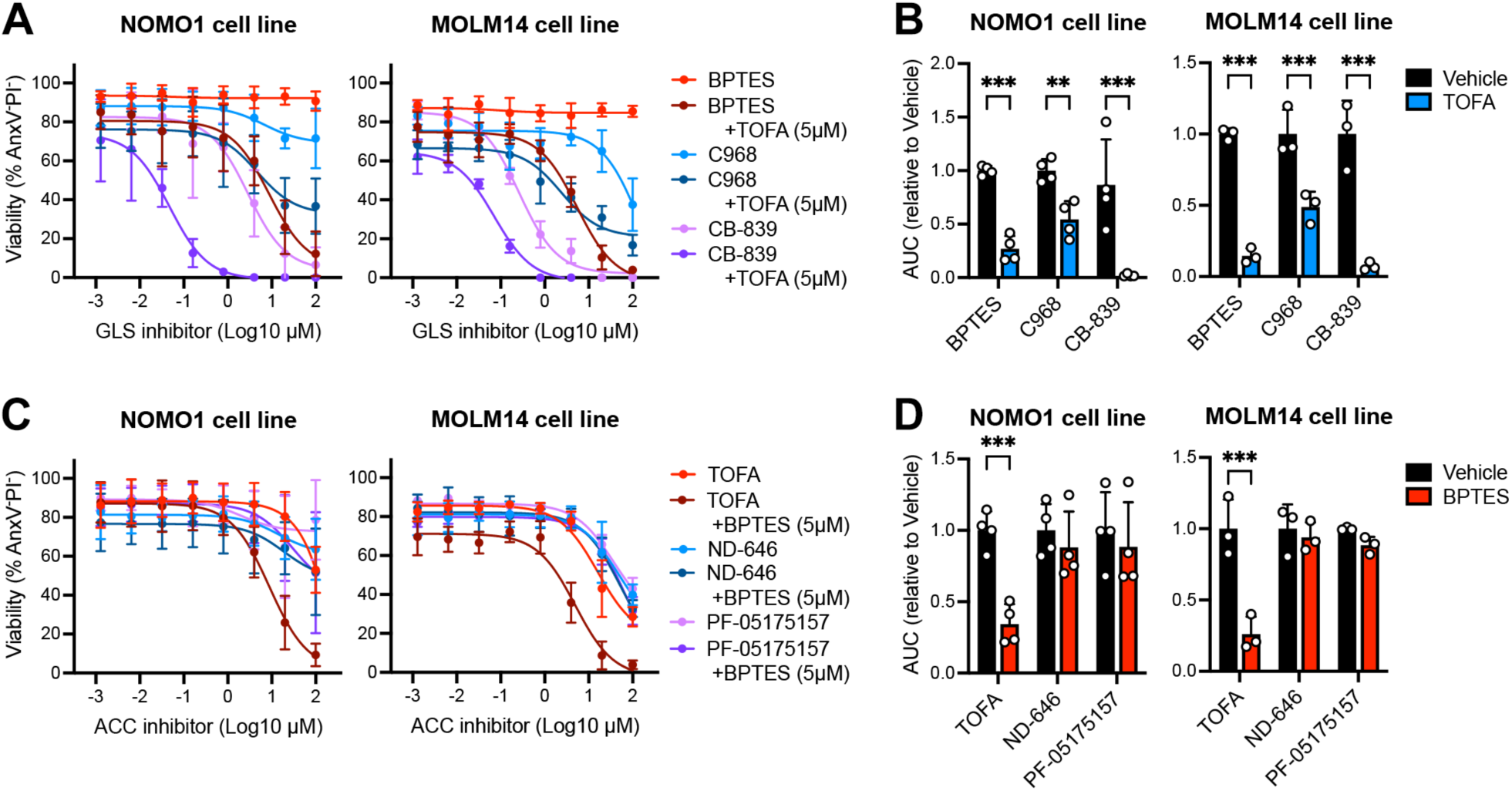
Cytotoxicity of TOFA is Unrelated to Inhibition of Lipogenesis. (**A**) Viability of AML cell lines (mean ± SD; N=4) treated for 72 hours with increasing doses of BPTES, C968 or CB-839 in the presence or absence of TOFA (5 µM). GLS, glutaminase. (**B**) Area under the curve (AUC) calculated from the viability data shown in A. (**C**) Viability of AML cell lines (mean ± SD; N=4) treated for 72 hours with increasing doses of TOFA, ND-646 or PF-05175157 in the presence or absence of BPTES (5 µM). ACC, acetyl-CoA carboxylase. (**D**) Area under the curve (AUC) calculated from the viability data shown in C. **p < 0.01, ***p < 0.001. See also Figure S4.

We then followed a similar approach for TOFA. Unexpectedly, the potent ACC inhibitors ND-646 and PF-05175157 did not affect AML cell viability or sensitivity to BPTES (Figure 4C-D), even though they efficiently inhibited *de novo* lipogenesis when used at a concentration of 5 µM (Figure S4A). Likewise, genetic knockdown of ACC1 did not induce NOMO1 cell death nor did it increase the response to BPTES (Figure S4B-C). TOFA alone or in combination with BPTES also had similar cytotoxic effects in cells lacking ACC1 compared to cells transduced with a control short hairpin RNA (shRNA), showing that ACC1 is not required for the action of TOFA (Figure S4C). Loss of ACC1 even slightly enhanced apoptosis induction by TOFA, although this effect was not significant for all shRNAs tested. Targeting *de novo* lipogenesis in an alternative way, by inhibiting fatty acid synthase (FASN) using the small molecules GSK2194069 or TVB-3166, also did not synergize with BPTES in inducing AML cell death (Figure S4A, Figure S4D). These results therefore show that the ability of TOFA to induce AML cell death and synergize with BPTES is unrelated to its inhibition of *de novo* lipogenesis.

### TOFA efficiently inhibits protein *S*-acylation

The structural similarity of TOFA to long chain fatty acids (Figure 5A) leads to its conversion into a CoA ester (TOFyl-CoA) after cellular uptake, which explains its ability to inhibit ACC enzymes^21^. Based on this knowledge, and our finding that the effects of TOFA in AML cells are unrelated to the inhibition of *de novo* lipogenesis or the increase in membrane lipid unsaturation but can be prevented by the addition of exogenous SFAs/MUFAs, we hypothesized that TOFA also inhibits other enzymes that use long chain fatty acyl-CoAs as substrate. Querying the Braunschweig Enzyme Database (BRENDA^30^) for enzymes that use palmitoyl-CoA or stearoyl-CoA as substrate led to a list of 21 enzyme classes relevant for mammalian cells (Table S4). To prioritize these potential targets, we retrieved genome-scale CRISPR screen data (DepMap Public 24Q2+Score, Chronos) and RNA-seq data (Batch-corrected TPM values) of AML cell lines (26 lines) from the DepMap portal^31^ for all enzymes potentially using long chain fatty acyl-CoAs as substrate. Plotting these data identified 10 enzymes that on average are highly expressed in AML cells lines and reduce their growth when lost (Figure 5B). The top hits were stearoyl-CoA desaturase (SCD), serine palmitoyltransferase long chain base subunit 1 and 2 (SPTLC1 and 2) and several zinc finger Asp-His-His-Cys motif-containing (zDHHC) *S*-acyltransferases (zDHHC8, 11, 18 and 20).

**Figure 5.**
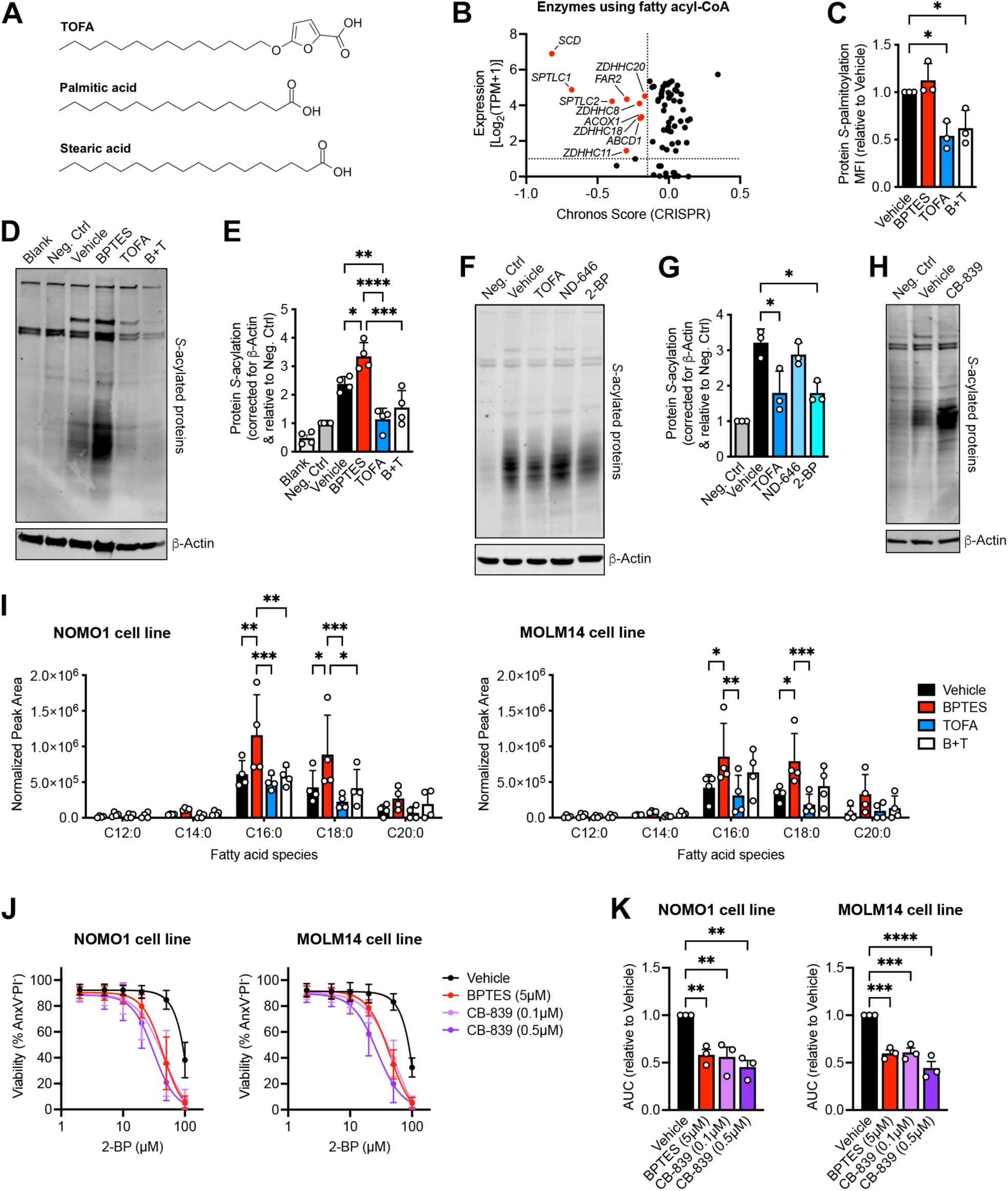
TOFA Inhibits Protein *S*-Acylation. (**A**) Schematic comparing the molecular structure of TOFA to long chain saturated fatty acids. (**B**) Gene expression versus dependency (from pooled CRISPR-Cas9 knockout screens; average of 26 human AML cell lines) for 70 enzymes using palmitoyl-CoA or stearoyl-CoA as substrate. (**C**) Protein *S*-palmitoylation revealed by click chemistry in NOMO1 cells treated for 24 hours with BPTES (5 µM) and/or TOFA (5 µM) in the presence of alkynyl-palmitate. MFI, mean fluorescence intensity. (**D-E**) Blot detection (D) and quantification (E) of total protein *S*-acylation through acyl-biotin exchange in NOMO1 AML cells treated for 24 hours with BPTES and/or TOFA. β-actin was used as loading control. (**F-G**) Blot detection (F) and quantification (G) of total protein *S*-acylation through acyl-biotin exchange in NOMO1 AML cells treated for 24 hours with TOFA (5 µM), ND-646 (5 µM), 2-BP (50 µM) or vehicle. β-actin was used as loading control. (**H**) Blot detection of total protein *S*-acylation through acyl-biotin exchange in NOMO1 AML cells treated for 24 hours with CB-839 (0.5 µM) or vehicle. β-actin was used as loading control. (**I**) Identification and quantification of fatty acid species covalently bound to proteins by thioester linkage in AML cell lines treated for 24 hours with BPTES and/or TOFA. (**J**) Viability of AML cell lines treated for 72 hours with increasing doses of 2-BP in the presence or absence of BPTES or CB-839. (**K**) Area under the curve (AUC) calculated from the viability data shown in J. *p < 0.05, **p < 0.01, ***p < 0.001, ****p < 0.0001. See also Figure S5.

We first focused on SCD, given that this enzyme showed the highest expression and largest susceptibility across AML cell lines (Figure 5B), was recently found to be a metabolic vulnerability of AML cells^32,33^, and has been identified as a potential target of TOFA^34^. SCD is a delta-9-desaturase that converts the saturated fatty acyl-CoAs palmitoyl-CoA and stearoyl-CoA into their monounsaturated equivalents, respectively palmitoleoyl-CoA and oleoyl-CoA. Since oleic acid was the most effective at preventing TOFA-induced cell death (Figure 3D), we hypothesized that AML cells may require MUFAs for a specific process unrelated to membrane saturation. In line with previous findings, we found that most AML cell lines were sensitive to SCD inhibitors (MF438, CAY10566, A9395572) at doses that effectively block enzyme activity (Figure S5A-C). However, apart from OCI-AML3 cells, treatment with SCD inhibitors did not sensitize AML cells to BPTES, showing that the effects of TOFA are likely not due to the inhibition of SCD (Figure S5B-C).

SPTLC1 and SPTLC2, both of which are highly expressed across AML cell lines and strongly reduced AML cell growth when lost, catalyze the first and rate-limiting step of ceramide biosynthesis^35^. Since we did not observe any change in the content of ceramides or other sphingolipids (glycosylceramides, sphingomyelins) of AML cells after TOFA treatment (Figure S3C), we deemed it unlikely that these enzymes are responsible for the effects of TOFA. Instead, we focused our attention on the zDHHC protein *S*-acyltransferases. Protein *S*-acylation is a reversible post-translational modification where fatty acids are covalently attached to proteins through a thioester linkage^36^. While *S*-acylation modifies proteins with long chain fatty acids between 14 and 22 carbons in length, it is often referred to as *S*-palmitoylation since palmitoyl-CoA is considered to be the most common fatty acid donor^36,37^. Through a click chemistry reaction using an azide-containing fluorescent dye on NOMO1 AML cells incubated with palmitic acid alkyne, we discovered that TOFA strongly reduces protein *S*-palmitoylation, while BPTES had little effect (Figure 5C). To better characterize protein *S*-acylation in AML cells we performed an acyl-biotin exchange assay in which all *S*-acylated proteins can be analyzed irrespective of the bound fatty acid moiety. This analysis confirmed that TOFA reduces protein *S*-acylation, but also showed an overall increase in NOMO1 AML cells treated with BPTES (Figure 5D-E). The effects of TOFA on protein *S*-acylation were comparable to those of 2-bromopalmitate (2-BP), a commonly used zDHHC inhibitor^38^, while the ACC inhibitor ND-646 did not alter protein *S*-acylation (Figure 5F-G), indicating that the ability of TOFA to inhibit protein *S*-acylation is independent of its effects on *de novo* lipogenesis. Like BPTES, treatment of NOMO1 AML cells with CB-839 increased overall protein *S*-acylation, showing that this is a common effect of GLS inhibition (Figure 5H).

Analysis of the fatty acid tails bound to AML cell proteins by GC-MS confirmed an overall increase in protein *S*-acylation with BPTES and decrease with TOFA, and further revealed that in addition to palmitic acid, stearic acid is commonly used for protein modification by AML cells (Figure 5I). We also detected arachidic acid (C20:0), especially in cells treated with BPTES, with minor contributions of other long chain SFAs. MUFAs or PUFAs were not found among the pool of fatty acids bound to proteins in AML cells. We also did not detect the methyl ester of TOFA in these analyses, indicating that TOFA itself is not covalently bound to proteins by a thioester linkage. The ability of TOFA to block protein *S*-acylation is thus likely due to its ability to occupy the fatty acid-binding pocket of the zDHHC enzymes, similar to 2-BP^38^. Combining 2-BP with the GLS inhibitors BPTES and CB-839 showed synergism in inducing AML cell death (Figure 5J-K), thus confirming that protein *S*-acylation is required for AML cells to metabolically adapt to the inhibition of glutaminolysis.

### Protein *S*-acylation preserves AML cell metabolic activity when glutaminolysis is inhibited

Protein *S*-acylation is thought to affect up to 20% of the human proteome, and as such it can regulate many cellular processes, including cell signaling, migration, division, communication, and metabolism^36^. To better understand the effects of BPTES and TOFA on this process, we analyzed *S*-acylated proteins in treated AML cells by performing an acyl-biotin exchange assay followed by streptavidin pull-down and proteomics analysis (Figure S6A). We identified a total of 382 proteins that were *S*-acylated in NOMO1 cells in at least one condition across two independent experiments (Table S5).

Unsupervised clustering of the top 100 proteins differing between groups, as determined by ANOVA, revealed 4 major clusters (Figure 6A, Table S6). A first set of proteins was enriched particularly in the vehicle condition (Class 1; 16 proteins), a second group was most abundant in BPTES-treated cells (Class 2; 61 proteins), a third cluster contained proteins that were unchanged by BPTES but lower in cells treated with TOFA or with the combination (Class 3; 19 proteins) and a very small fourth group (Class 4; 4 proteins) contained proteins that were enriched in cells treated with TOFA or with the combination of BPTES and TOFA.

**Figure 6.**
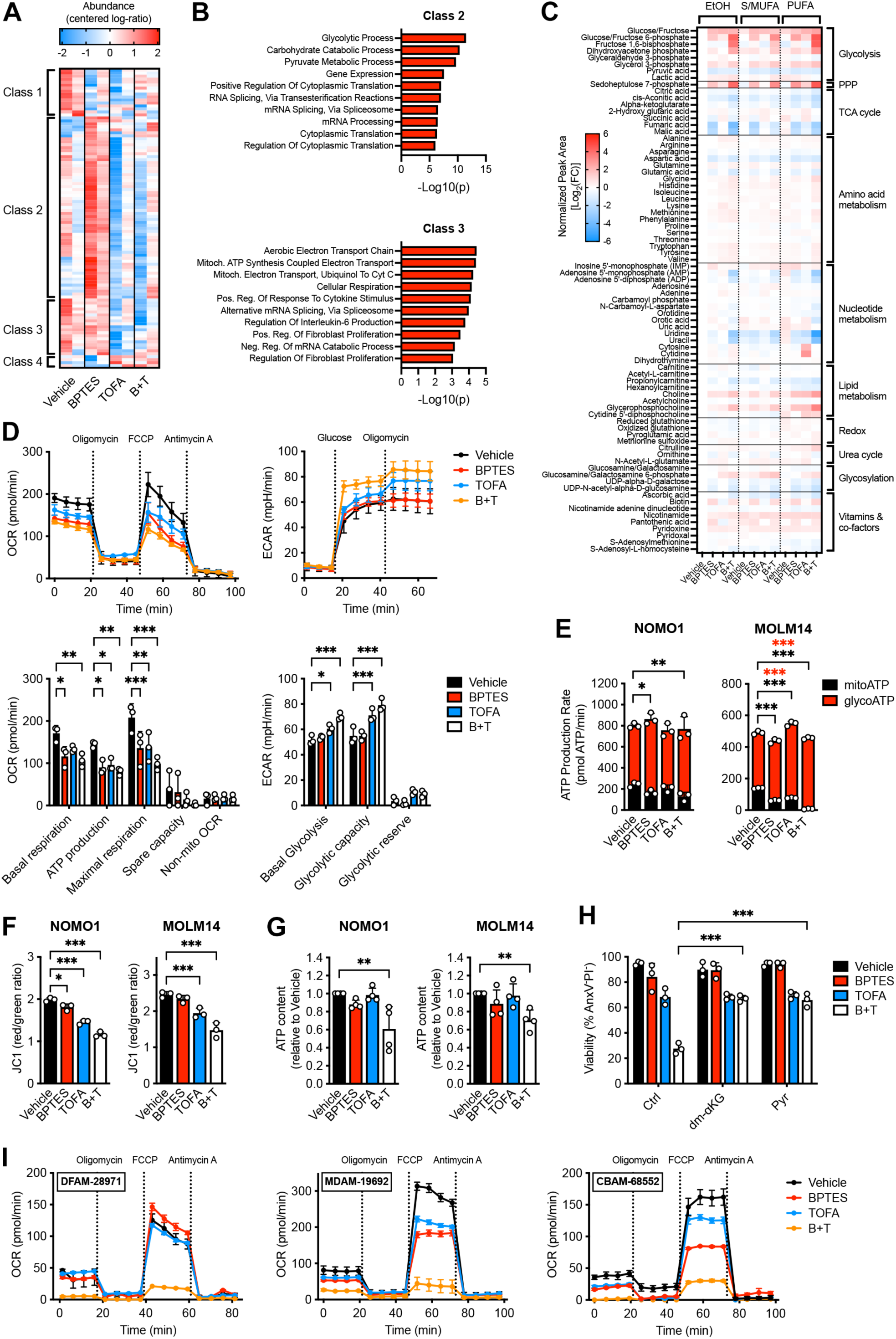
Protein *S*-Acylation Supports Metabolic Plasticity in AML (legend on next page) (**A**) Heatmap of proteomics data showing unsupervised clustering of the top 100 proteins differently *S*-acylated between NOMO1 AML cells treated for 24 hours with BPTES (5 µM) and/or TOFA (5 µM), as determined by ANOVA. Four Classes of proteins showing different *S*-acylation profiles between groups were identified through manual data analysis. (**B**) Enrichment analysis of proteins included in Class 2 and 3 identified from the proteomics data shown in A. (**C**) Heatmap of the metabolic profile of NOMO1 AML cells treated for 24 hours with BPTES and/or TOFA. (**D**) Real-time oxygen consumption rate (OCR; top left) and extracellular acidification rate (ECAR; top right) of NOMO1 AML cells (mean ± SD; N=3) treated for 24 hours with BPTES and/or TOFA, measured by Seahorse assay. Graphs show quantification of different OXPHOS parameters from OCR data (bottom left) and glycolysis parameters from ECAR data (bottom right). (**E**) Mitochondrial ATP production rate (mitoATP) and glycolytic ATP production rate (glycoATP) of AML cell lines treated for 24 hours with BPTES and/or TOFA, measured by Seahorse assay. (**F-G**) Mitochondrial membrane potential measured by JC1 staining (F), and intracellular levels of ATP measured by CellTiter-Glo assay (G) in AML cell lines treated for 24 hours with BPTES and/or TOFA. (**H**) Viability of NOMO1 AML cells treated for 72 hours with BPTES and/or TOFA in the presence or absence of dimethyl alpha-ketoglutarate (dm-αKG; 5 mM) or sodium pyruvate (Pyr; 1 mM). (**I**) Real-time OCR of three different AML PDX lines (mean ± SD; N=3) treated for 24 hours with BPTES and/or TOFA, measured by Seahorse assay. *p < 0.05, **p < 0.01, ***p < 0.001. See also Figure S6.

We first focused on Class 2, containing proteins that show increased *S*-acylation in response to BPTES, which is blocked in the presence of TOFA. We hypothesized that proteins in this Class may allow AML cells to survive the inhibition of glutaminolysis by undergoing increased *S*-acylation, explaining the synergism between BPTES and TOFA. Enrichment analysis showed that Class 2 was highly enriched for proteins involved in glycolysis (Figure 6B). Many glycolytic enzymes showed increased *S*-acylation in response to BPTES, including glucose-6-phosphate isomerase (GPI), aldolase (ALDOA, ALDOC), glyceraldehyde 3-phosphate dehydrogenase (GAPDH), phosphoglycerate kinase (PGK1), enolase (ENO1), pyruvate kinase (PKM) and lactate dehydrogenase (LDHA) (Figure S6B). Western Blot analysis on pulled-down proteins after acyl-biotin exchange validated these findings, showing that GAPDH and PKM *S*-acylation increased with BPTES and was inhibited by TOFA, while the glycolytic enzyme hexokinase 2 (HK2), which was not picked-up by proteomics, was confirmed to only have a very low degree of *S*-acylation (Figure S6C). Besides glycolytic enzymes, Class 2 was enriched for proteins implicated in RNA processing and splicing, translation, and also contained several mitochondrial proteins (Figure S6B), including voltage dependent anion channel 2 (VDAC2), electron transport chain subunits and malate dehydrogenase 2 (MDH2) that was previously found to be *S*-acylated by zDHHC18 in ovarian cancer cells in response to glutamine starvation^39^. Increased *S*-acylation of MDH2 in response to BPTES was confirmed by Western Blot (Figure S6C).

While Class 1 and Class 4 did not show enrichment for any specific protein types or pathways, Class 3, containing proteins that do not show clear changes in *S*-acylation in response to BPTES but much lower *S*-acylation upon TOFA treatment, was also enriched for proteins involved in mitochondrial metabolism (Figure 6B), including TOMM20, cytochrome C and NDUFB10 (Figure S6B, Table S6). Adenylate kinase 2 (AK2), which was recently shown to be *S*-acylated by zDHHC21 in AML cells to support OXPHOS^40^, was not included in the top 100 proteins differing between groups but showed changes in response to BPTES and TOFA comparable to the other mitochondrial proteins (Figure S6B-C).

Given the *S*-acylation changes induced by BPTES and TOFA in AML cells, we decided to examine glycolytic and mitochondrial metabolism in more detail. We first performed metabolomics analysis, which showed that while BPTES moderately increased the level of most glycolytic metabolites and TOFA alone had little effect, the combination of BPTES and TOFA induced a large increase in upstream glycolytic metabolites (hexose monophosphate, fructose bisphosphate, dihydroxyacetone phosphate) as well as the pentose phosphate pathway intermediate sedoheptulose 7-phosphate (Figure 6C, Table S7). In contrast, the glycolytic end products pyruvate and lactate showed only small changes. In line with our prior tracing analysis (Figure 2C, Figure S2B), most TCA cycle metabolites were depleted in NOMO1 cells treated with BPTES, and this effect was exacerbated in combination with TOFA (Figure 6C, Table S7). Similar profiles were observed for metabolites in pathways linked to mitochondrial activity, including nucleotide metabolism intermediates (AMP, ADP, orotate, uridine, uracil) and non-essential amino acids such as aspartate and glutamate. In line with their effects on cell viability (Figure 3C), we found that supplementation of the culture medium with a mixture of SFAs/MUFAs, but not with PUFAs, partially restored the levels of both glycolytic and TCA cycle intermediates in NOMO1 AML cells treated with the combination of BPTES and TOFA (Figure 6C, Table S7).

In-depth analysis of mitochondrial activity showed that NOMO1 and MOLM14 AML cells treated with either BPTES or TOFA had a decreased oxygen consumption rate (OCR) indicative of reduced OXPHOS activity, while the combination of BPTES and TOFA further reduced the OCR (Figure 6D, Figure S6D). In line with this, BPTES and TOFA synergistically decreased mitochondrial ATP production (Figure 6E) as well as the mitochondrial membrane potential of AML cells (Figure 6F), while mitochondrial mass was not affected (Figure S6E). For cells treated with TOFA alone or in combination with BPTES, the drop in OXPHOS activity was mirrored by an increase in the extracellular acidification rate (ECAR), a measure for glycolytic activity (Figure 6D, Figure S6D). This compensation was also reflected in the glycolytic ATP production rate (Figure 6E) and the glucose uptake rate (Figure S6F). Surprisingly, cells treated with BPTES did not show evident changes in their ECAR, glycolytic ATP production or glucose uptake, even though BPTES changed the *S*-acylation profile of many glycolytic enzymes in NOMO1 cells, as described above. In addition, despite compensatory changes in glycolysis, AML cells treated with the combination of BPTES and TOFA had significantly lower cellular ATP levels (Figure 6G). These results indicate that the mitochondrial effects of BPTES and TOFA are primarily responsible for their synergism. Accordingly, rescue experiments with the anaplerotic substrates alpha-ketoglutarate and pyruvate restored viability of NOMO1 cells treated with the combination of BPTES and TOFA (Figure 6H). Together, these data prove that TOFA, by inhibiting the *S*-acylation of several mitochondrial proteins involved in the TCA cycle and in cellular respiration, prevents metabolic plasticity and renders AML cells unable to compensate for reduced glutaminolysis.

To further validate these findings, we used primary AML patient samples maintained as patient-derived xenografts (PDX) in immunocompromised mice, which are more amenable to *ex vivo* metabolic analysis compared to fresh patient-derived samples. For all three tested models (Table S1) we could confirm a more pronounced induction of cell death by the combination of BPTES and TOFA compared to either metabolic inhibitor alone (Figure S6G). Single treatment of AML-PDX cells with either BPTES or TOFA reduced the OCR in two out of three tested lines, but the combination strongly reduced mitochondrial OCR and maximal respiratory capacity across all tested models (Figure 6I, Figure S6H). These results confirm that like in AML cell lines, the combination of BPTES and TOFA prevents metabolic adaptation and triggers a mitochondrial collapse in AML patient-derived cells.

### Healthy HSPCs do not depend on protein *S*-acylation for metabolic adaptation to GLS inhibition

In contrast to AML cells, non-malignant HSPCs derived from cord blood or bone marrow do not undergo apoptosis in response to BPTES and TOFA treatment (Figure 1), even though the effects of BPTES on glutaminolysis are very similar to those observed in AML cells (Figure 2C, Figure S2B). Compared to AML cells, CB-CD34^+^ cells showed a lower degree of protein *S*-palmitoylation and overall protein *S*-acylation (Figure 7A-C), and treatment with BPTES or TOFA did not significantly alter protein *S*-acylation (Figure 7D-E). In addition, neither BPTES, nor TOFA, nor the combination significantly lowered the mitochondrial membrane potential of CB-CD34^+^ cells (Figure 7F), suggesting that HSPCs possess other mechanisms to compensate for the effects of BPTES.

**Figure 7.**
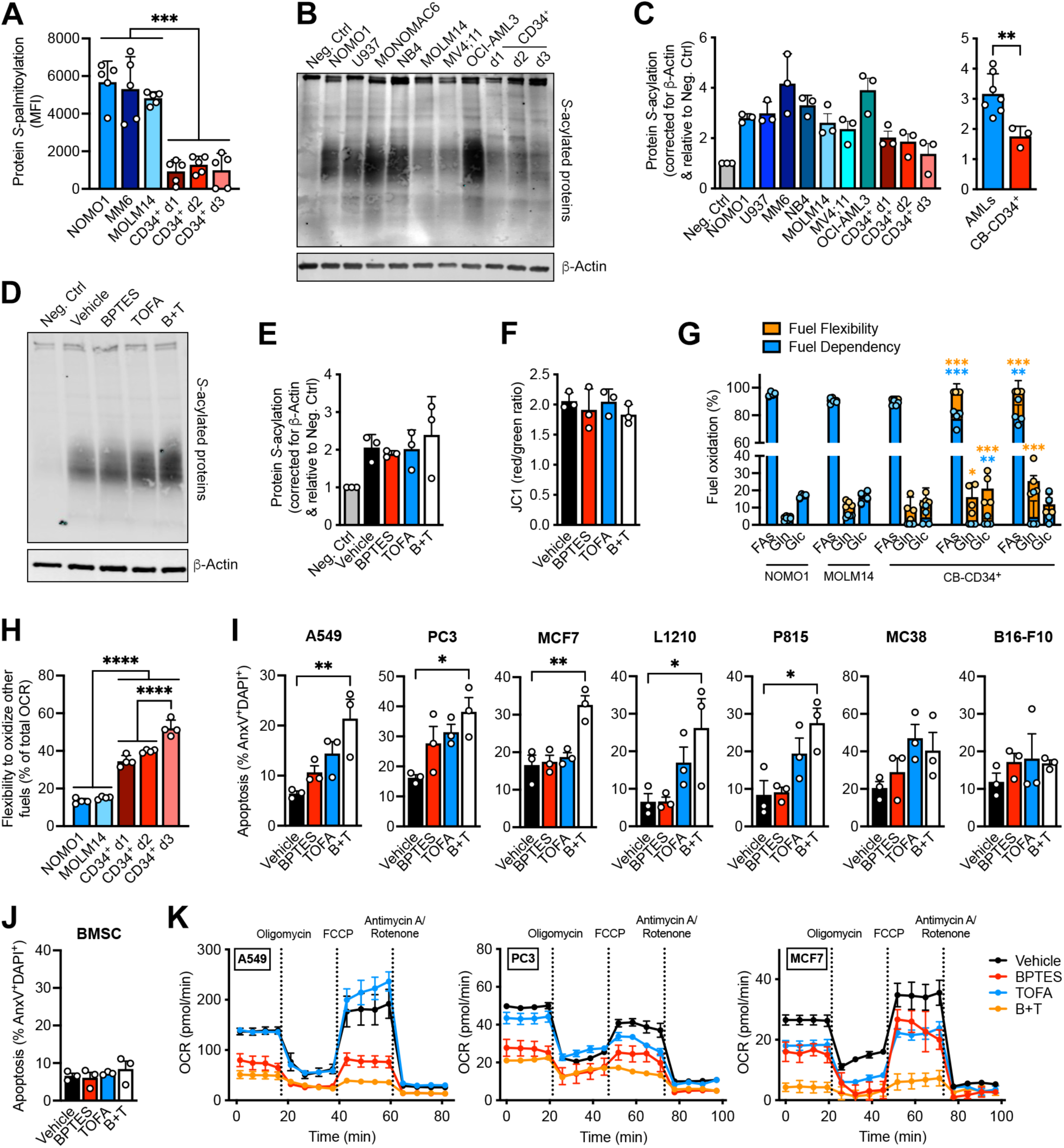
Differential Regulation of Metabolic Adaptation in Cancer and Healthy Cells. (**A**) Protein *S*-palmitoylation revealed by click chemistry in AML cell lines or cord blood-derived CD34^+^ cells (three independent donors shown), cultured for 24 hours in the presence of alkynyl-palmitate. MFI, mean fluorescence intensity. (**B-C**) Blot detection (B) and quantification (C) of total protein *S*-acylation through acyl-biotin exchange in AML cell lines or cord blood-derived CD34^+^ cells. β-actin was used as loading control. (**D-E**) Blot detection (D) and quantification (E) of total protein *S*-acylation through acyl-biotin exchange in cord blood-derived CD34^+^ cells treated for 24 hours with BPTES (5 µM) and/or TOFA (5 µM). β-actin was used as loading control. (**F**) Mitochondrial membrane potential measured by JC1 staining in cord blood-derived CD34^+^ cells treated for 24 hours with BPTES and/or TOFA. (**G**) Substrate flexibility and dependency for fueling OXPHOS in AML cell lines or cord blood-derived CD34^+^ cells, measured by Seahorse assay. FAs, fatty acids; Gln, glutamine; Glc, glucose. (**H**) Ability to use substrates other than glucose, glutamine or fatty acids to fuel OXPHOS in AML cell lines or cord blood-derived CD34^+^ cells, measured by Seahorse assay. (**I-J**) Apoptosis of different human and mouse cancer cell lines (I) or healthy human bone marrow stromal cells (BMSC; J) treated for 72 hours with BPTES and/or TOFA. (**K**) Real-time oxygen consumption rate (OCR) of three different human solid tumor cell lines (mean ± SD; N=3) treated for 24 hours with BPTES and/or TOFA, measured by Seahorse assay. *p < 0.05, **p < 0.01, ***p < 0.001, ****p < 0.0001. See also Figure S7.

To better understand the difference in metabolic adaptability between AML cells and HSPCs, we measured changes in OCR in response to different combinations of BPTES (inhibiting glutamine oxidation), the mitochondrial pyruvate carrier inhibitor UK5099 (inhibiting glucose/pyruvate oxidation) and the carnitine palmitoyltransferase I inhibitor etomoxir (inhibiting fatty acid oxidation), and calculated fuel dependency and flexibility (Figure 7G). Compared to AML cell lines, CB-CD34^+^ cells were much less dependent on individual substrates and showed higher flexibility to switch between glutamine, glucose and fatty acid oxidation to support OXPHOS. In addition, non-malignant HSPCs were able to maintain 40-50% of their mitochondrial OCR even when glutamine, glucose/pyruvate and fatty acid oxidation were inhibited simultaneously, indicating their ability to oxidize additional substrates (Figure 7H). In line with this, there was no reduction in viability when CB-CD34^+^ cells were treated with etomoxir in combination with BPTES and TOFA (Figure S7A). Together, these results show that HSPCs have a much higher mitochondrial flexibility compared to AML cells, allowing them to readily compensate for the effects of BPTES without changing overall protein *S*-acylation, and thus avoid a metabolic collapse in the presence of TOFA.

### BPTES and TOFA induce apoptosis and mitochondrial dysfunction in other cancer types

The combination of BPTES and TOFA induces a metabolic collapse in AML cells but spares normal HSPCs due to intrinsic differences in metabolic flexibility. With this in mind, we investigated whether other tumor types would also be sensitive to the combination of BPTES and TOFA by screening several human and mouse cancer cell lines. For some cell lines, including MCF7 human breast cancer cells, L1210 mouse lymphocytic leukemia cells and P815 mouse mastocytoma cells, a synergistic induction of apoptosis was observed when BPTES and TOFA were combined (Figure 7I, Figure S7B). In contrast, A549 human lung cancer cells and PC3 human prostate cancer cells showed additive effects of both compounds, while MC38 mouse colon cancer cells and B16-F10 mouse melanoma cells did not undergo apoptosis when exposed to BPTES and/or TOFA at the concentrations tested (Figure 7I, Figure S7B). Healthy human bone marrow stromal cells (BMSCs), like HSPCs, did not show any signs of apoptosis when treated with either BPTES, TOFA or the combination (Figure 7J). Irrespective of the induction of apoptosis, all tested cancer cell lines showed a lower number of viable cells in the cultures after 72 hours of treatment with BPTES and TOFA compared to the single compounds (Figure S7C). This was less pronounced with BMSCs, although TOFA on its own did reduce their growth (Figure S7D). We then examined the impact of BPTES and/or TOFA on mitochondrial metabolism in A549, PC3 and MCF7 cells. In MCF7 cells, treatment with either BPTES or TOFA alone reduced the OCR, while in A549 and PC3 cells only BPTES had a significant effect (Figure 7K, Figure S7E). In all 3 cell lines the combination of BPTES and TOFA showed a stronger reduction of the OCR, although this effect was the most pronounced in MCF7 cells, in line with the synergistic induction of cell death in this cell line (Figure 7I, Figure S7B). Thus, our results indicate that other cancer types may also depend on protein *S*-acylation to metabolically adapt to the inhibition of glutaminolysis, while this is not seen in non-cancerous cells.

## Discussion

In the century following Otto Warburg’s discovery that carcinoma cells exhibit increased glycolysis compared to normal epithelial cells^41^, numerous studies have contributed to our understanding of the distinct metabolic features and dependencies of solid tumor and hematological cancer cells^42^. However, translating these insights into new therapies has proven challenging. The reasons for this are still incompletely understood, but the ability of cancer cells to rewire their metabolism in response to inhibition of a single pathway has emerged as a key feature^16^. In the current study, we tackled this challenge by exposing AML cells to random combinations of metabolic inhibitors with the goal of identifying strategies to overcome metabolic adaptation upon single pathway inhibition. Our results show that AML cells, including LSCs, can be sensitized to GLS inhibitors by concurrent treatment with TOFA. We discovered that TOFA exerts this effect through a non-canonical inhibition of protein *S*-acyltransferases, thus preventing a shift in metabolic enzyme *S*-acylation that confers metabolic plasticity to AML cells. The hyper-dependency on protein *S*-acylation dynamics in AML cells appears to stem from a loss of the inherent metabolic flexibility that allows healthy hematopoietic progenitors to readily switch between nutrients to fuel OXPHOS. These findings not only provide new biological and pharmacological insights but also suggest an alternative therapeutic approach to potentiate metabolic cancer therapies such as GLS inhibitors.

When considering metabolic adaptability in cancer, glutaminolysis may be the pathway that exemplifies this phenomenon better than any other. Many cancer types show glutaminolysis hyperactivation and some have even been termed “glutamine addicted”^43^, yet GLS inhibitors such as CB-839 have underperformed in the clinic despite good safety profiles and tolerability, with cancer cells in non-responders showing evidence of metabolic compensation^44–46^. Our results show that upon GLS inhibition, AML cells undergo a shift in protein *S*-acylation to rewire their metabolism and avoid cell death induction, even though proliferation is reduced. One of the modified pathways is glycolysis, with several glycolytic enzymes showing a stark increase in *S*-acylation after GLS inhibition. This is reminiscent of previous findings in MYC-driven lung tumors and pancreatic as well as colorectal cancer cell lines, where inhibition of glutamine catabolism led to a compensatory increase in glycolytic activity^47,48^, while in AML it has been shown that interference with glycolysis increases the dependence on glutaminolysis^49^. However, in our current experiments the overall increase in glycolytic activity with BPTES was relatively mild and we only observed clear changes when AML cells were concomitantly treated with BPTES and TOFA, and upstream glycolytic metabolites accumulated massively. These data suggest that the increased *S*-acylation of glycolytic enzymes does not simply promote a glycolytic switch, further confirmed by the fact that TOFA did not reduce, but rather increased, glycolytic activity and that our initial combinatorial screening failed to show strong synergies between BPTES and compounds targeting glucose metabolism (TEPP46, FX11, DCA). Several glycolytic enzymes have previously been shown to be *S*-acylated, but while *S*-acylation of GLUT1 or LDHA increases glycolysis^50,51^, *S*-acylation of GAPDH or PKM2 was shown to reduce glycolytic flux^52,53^. It is therefore likely that the general increase in glycolytic enzyme *S*-acylation in BPTES-treated AML cells does not increase pathway activity *per se* but influences glycolysis in another way. Given that *S*-acylation is a key for mediating protein membrane targeting and compartmentalization in the cell^36^, it is conceivable that increased *S*-acylation of glycolytic enzymes alters their location in the cell and could for example tether them to the mitochondrial membrane to enhance metabolic coupling between glycolysis and OXPHOS^54–56^ or even directly impact mitochondrial apoptotic signaling^57^.

In this regard, it is of interest to note that the metabolic synergy between BPTES and TOFA occurs primarily at the level of mitochondrial respiration in AML cells. *S*-acylation of MDH2, which increases in response to BPTES and is inhibited by TOFA, may play an important role in this effect. *S*-palmitoylation of MDH2 was previously shown to support OXPHOS in ovarian cancer cells, an effect further enhanced by glutamine deprivation^39^. Exactly how increased MDH2 *S*-acylation allows cancer cells to survive reduced glutamine catabolism is still unclear. Given that this enzyme is part of both the TCA cycle and the malate-aspartate shuttle, its post-translational modification may impact the relative involvement of MDH2 in both pathways by stabilizing specific metabolic enzyme complexes, as previously suggested by Kostiuk *et al.*^58^. In addition to MDH2, AML cells show changes in the *S*-acylation of several other mitochondrial proteins in response to BPTES, but whether and how these changes help to maintain mitochondrial activity in AML cells when glutaminolysis is reduced remains to be determined. Other mitochondrial enzymes did not undergo changes in *S*-acylation in response to BPTES but were strongly affected by TOFA, such as AK2. *S*-acylation of AK2 by ZDHHC21 was shown to be key for maintaining mitochondrial activity in AML cells^40^, and while we did not observe changes in *S*-acylation of AK2 in response to BPTES, the activity of this enzyme may become particularly essential in AML cells when TCA cycle activity is reduced due to glutaminolysis inhibition. Together, our findings reveal that while AML cells possess the metabolic plasticity to survive inhibition of glutaminolysis or inhibition of *S*-acylation when occurring independently, the combination of both is the proverbial straw that breaks the camel’s back and induces a complete collapse of mitochondrial respiration.

TOFA was originally developed as a hypolipidemic agent able to lower serum cholesterol and triglycerides in rats^59^ and was later discovered to exert these effects through the inhibition of ACC by TOFyl-CoA^21^. A subsequent study showed that in addition to ACC, TOFyl-CoA also inhibits SCD1^34^. We found that in AML cells, neither inhibition of ACC or SCD1 can explain the cytotoxic effects of TOFA and its synergism with GLS inhibitors. Treatment with different unrelated ACC inhibitors or genetic knockdown of ACC1 did not affect AML cell viability, in line with previous reports showing that ACC1 may actually have a tumor suppressor role in AML^60,61^. While SCD1 does represent a metabolic vulnerability of AML cells, as previously shown^32,33^ and confirmed by our experiments, we did not observe a synergism with GLS inhibitors, unlike TOFA. This made us hypothesize that TOFA has an additional target in AML cells. Using a combination of available gene expression and CRISPR screening data and our own lipidomics and fatty acid rescue experiments we narrowed down the list of potential targets to the zDHHC protein *S*-acyltransferases and confirmed that TOFA efficiently blocks protein *S*-acylation. Further studies will be required to fully elucidate the mechanism through which TOFA inhibits protein *S*-acylation, including binding studies to see if TOFA binds to and prevents the catalytic activity of the zDHHC enzymes, if it does so by occupying the fatty acid-binding pocket of these enzymes as shown for 2-BP^38^ and if TOFA has any selectivity among the 23 zDHHC family members. Our findings also warrant a reevaluation of the conclusions of previous studies that used TOFA as a sole agent to inhibit ACC enzymes or *de novo* lipogenesis and underscore the importance for future studies using TOFA to consider the effects of this molecule on protein *S*-acylation.

Our study also sheds additional light on fatty acid metabolism in AML. While AML cells exhibit a higher rate of *de novo* lipogenesis compared to normal HSPCs, it is overall still relatively limited (8-10% of the total cellular palmitate pool is synthetized *de novo* in 24 hours) and AML cells do not depend on this pathway to maintain viability. This indicates that unlike some other cancer types^62^, the dependence on *de novo* lipogenesis is limited in AML. In contrast, AML cells readily take up exogenous fatty acids irrespective of their saturation degree, tolerate a high degree of lipid unsaturation in their membranes and compensate for a reduction in fatty acid synthesis by taking up even more exogenous fatty acids. In line with this, we found that most AML cell lines are sensitive to exogenous lipid deprivation, which further sensitizes them to TOFA (data not shown). However, while AML cells show flexibility in the source of fatty acids, they do depend on certain lipogenic enzymes such as SCD1 to maintain the optimal balance between individual fatty acid species. AML cells may particularly depend on MUFAs such as oleic acid, which we found to be most effective at rescuing the effects of TOFA when supplemented exogenously as a single fatty acid. Given that we only detected long chain SFAs including palmitic acid, stearic acid and arachidic acid to be covalently bound to proteins in AML cells, and not MUFAs or PUFAs, oleic acid may prevent the effects of TOFA by directly replenishing the MUFA pool, thus reducing the depletion of SFAs by SCD1 and freeing up the CoA esters of these FAs to compete with TOFyl-CoA for binding to zDHHC enzymes. The accuracy of this proposed mechanism remains to be determined.

Targeting the increased mitochondrial respiration of AML blasts and LSCs has been pursued extensively but clinical success remains limited, mainly due to toxicity issues and metabolic compensation. Targeting a single pathway fueling OXPHOS, such as glutaminolysis, shows limited efficacy. Could a combination of GLS inhibitors with TOFA overcome these limitations? We observed inherent differences between normal and cancer cells in the way they metabolically compensate for a loss of glutaminolysis activity, indicating that there may be a therapeutic window. The ability of normal cells to more readily switch between nutrients fueling the TCA cycle without depending on concurrent changes in protein *S*-acylation suggests that TOFA may present a way to specifically target the aberrant mitochondrial metabolism of AML and other cancer cells that does not induce a full shutdown in healthy tissues. Yet, several hurdles remain. TOFA does not have good drug-like properties, and its bioavailability *in vivo* is unclear. The fact that it has several targets will also complicate clinical translation. Our current findings, along with other recent studies, do underscore the importance of protein *S*-acylation for mitochondrial activity and metabolic plasticity in cancer cells and encourage additional studies to further elucidate the role and therapeutic potential of this metabolic process.

### Limitations of the study

While we identify protein *S*-acyltransferases as a new target of TOFA, we cannot exclude that its ability to block ACC and SCD1 (and potentially also other enzymes using acyl-CoA as substrate) contributes to the cytotoxic effects of this drug on AML cells and its synergism with BPTES. 2-BP, which also resembles a long-chain saturated fatty acid, may have similar pleotropic effects. Future studies using genetic targeting of zDHHC enzymes in combination with GLS inhibition can provide more clarity on this aspect. A second limitation of this study is that it is not currently feasible to characterize the synthetic lethality of TOFA and GLS inhibitors on AML cells *in vivo*. While several excellent, orally available GLS inhibitors have been developed, including CB-839, TOFA is much more challenging to administer to mice due to its very low aqueous solubility. Genetic targeting of zDHHCs could again be an alternative but may prove challenging given that this enzyme family has 23 members and that it is not yet known which member(s) need to be inhibited to achieve synergy with GLS inhibition.

## Acknowledgements

We thank Guido Bommer from the Biochemistry and Metabolic Research Group at de Duve Institute for access to and help with GC-MS analysis; Antony Letai (Dana-Farber Cancer Institute, Harvard Medical School) for help in designing the BH3 profiling assays; Nicolas Dauguet from the CYTF platform at the de Duve Institute and Joyce LaVecchio and Nema Kheradmand from the Flow Cytometry Core at the Harvard Department of Stem Cell and Regenerative Biology for help with flow cytometry analysis and cell sorting; Gaëtan Herinckx from the MASSPROT platform at de Duve Institute for help with sample preparation for proteomics analysis; the M.D. Anderson Cancer Center and Center for Patient-Derived Models at DFCI for the PDX models; and all members of the van Gastel and Scadden laboratories for helpful discussions. This work was supported by the Belgian Foundation Against Cancer [F/2020/1440, ATE-2022/1863, F/2024/2556 to N.v.G.], the Fund for Scientific Research - FNRS [F.R.S.-FNRS; A5/5-CQ/135, M4/1/2/5-MIS/BEJ to N.v.G.], Alex’s Lemonade Stand Foundation and Tap Cancer Out (Young Investigator Award to N.v.G.). N.B. was supported by a Chargée de Recherches postdoctoral fellowship from the F.R.S.-FNRS [A4/5-MCF/DEA-CR27]. A.E. was supported by a Rubicon postdoctoral research fellowship from the Netherlands Organisation for Health Research and Development (ZonMw) - Netherlands Organisation for Scientific Research (NWO) [2022/14232/ZONMW] and a Chargée de Recherche postdoctoral fellowship from the F.R.S.-FNRS [A4/5-CR179]. V.W.D. was supported by a postdoctoral fellowship from the Terri Brodeur Breast Cancer Foundation. M.A.K. and G.S. were funded by NIH grants 1R01DK075850-01 and 1R01CA160458-01A. A.A.L. is a Scholar of Blood Cancer United (formerly the Leukemia & Lymphoma Society). The MASSPROT platform (D.V.) was supported by the Baillet Latour Fund. D.T.S. was supported by the Harvard Stem Cell Institute and the Ludwig Center at Harvard, as well as the Gerald and Darlene Jordan Chair of Medicine at Harvard. The funders had no role in study design, data collection and analysis, decision to publish, or preparation of the manuscript. Schematics in Figure 1F and Figure S6A were created with BioRender.com.

## Author Contributions

N.v.G. and D.T.S. conceived the project and initiated the study. N.B., A.E., V.W.D., D.T.S. and N.v.G. designed the experiments, and interpreted results. N.B., A.E., A.S., V.W.D., P.L.C., F.L., Y.L. and N.v.G. performed the *in vitro* experiments and analyzed the data. V.W.D. designed, executed and interpreted BH3 profiling experiments. M.A.K. and G.S. designed, executed (M.A.K.) and interpreted stable isotope tracing experiments. C.V. and S.A.T. designed, executed and interpreted lipidomics and metabolomics analysis. A.A.L., H.C. and M.L. provided and analyzed patient samples and coordinated ethical approvals. D.V. designed, executed and interpreted proteomics analysis. N.B., A.E. and N.v.G. wrote and edited the manuscript with help from coauthors. N.v.G. provided overall project leadership. All authors revised the manuscript and approved its content.

## STAR Methods

## KEY RESOURCES TABLE

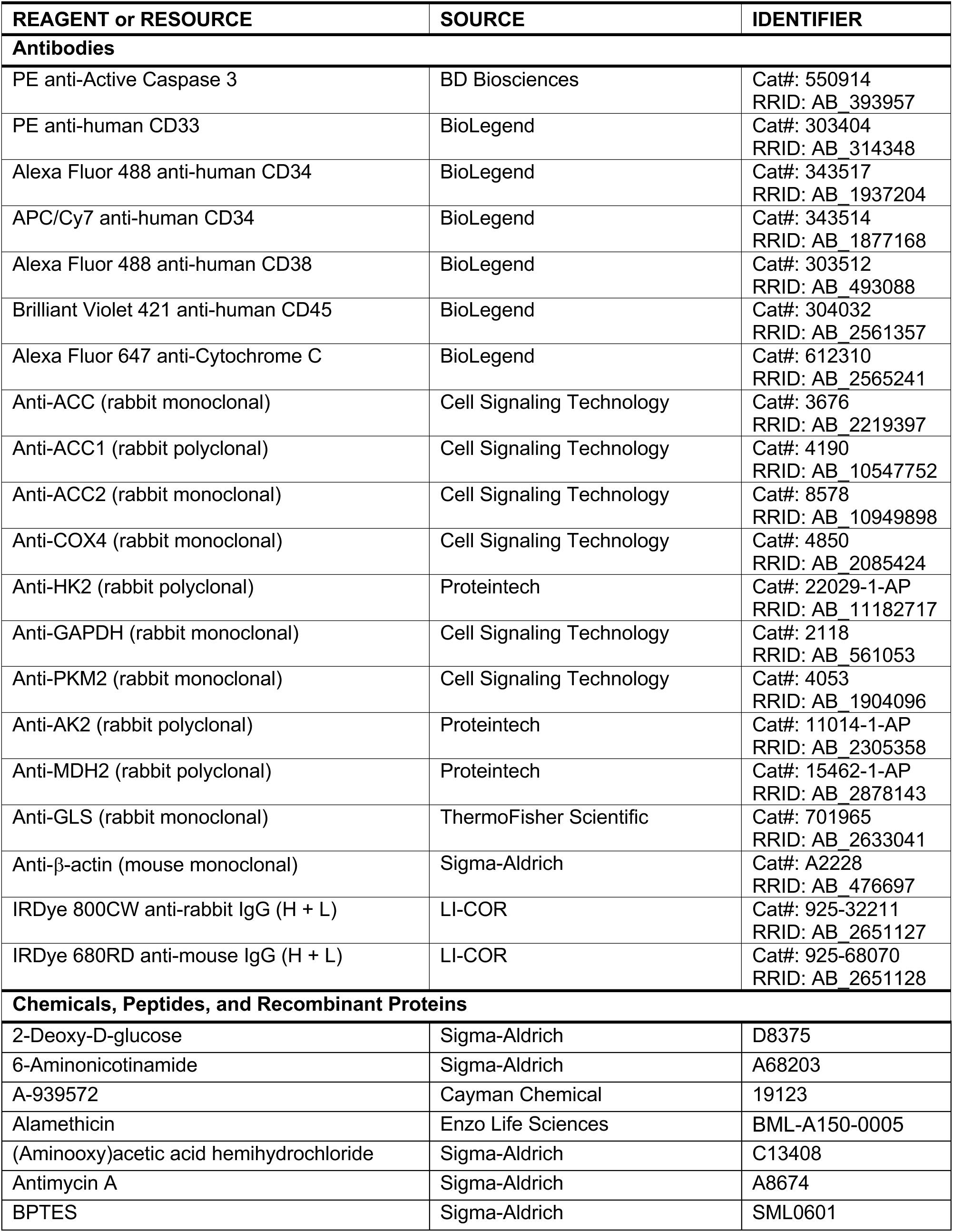
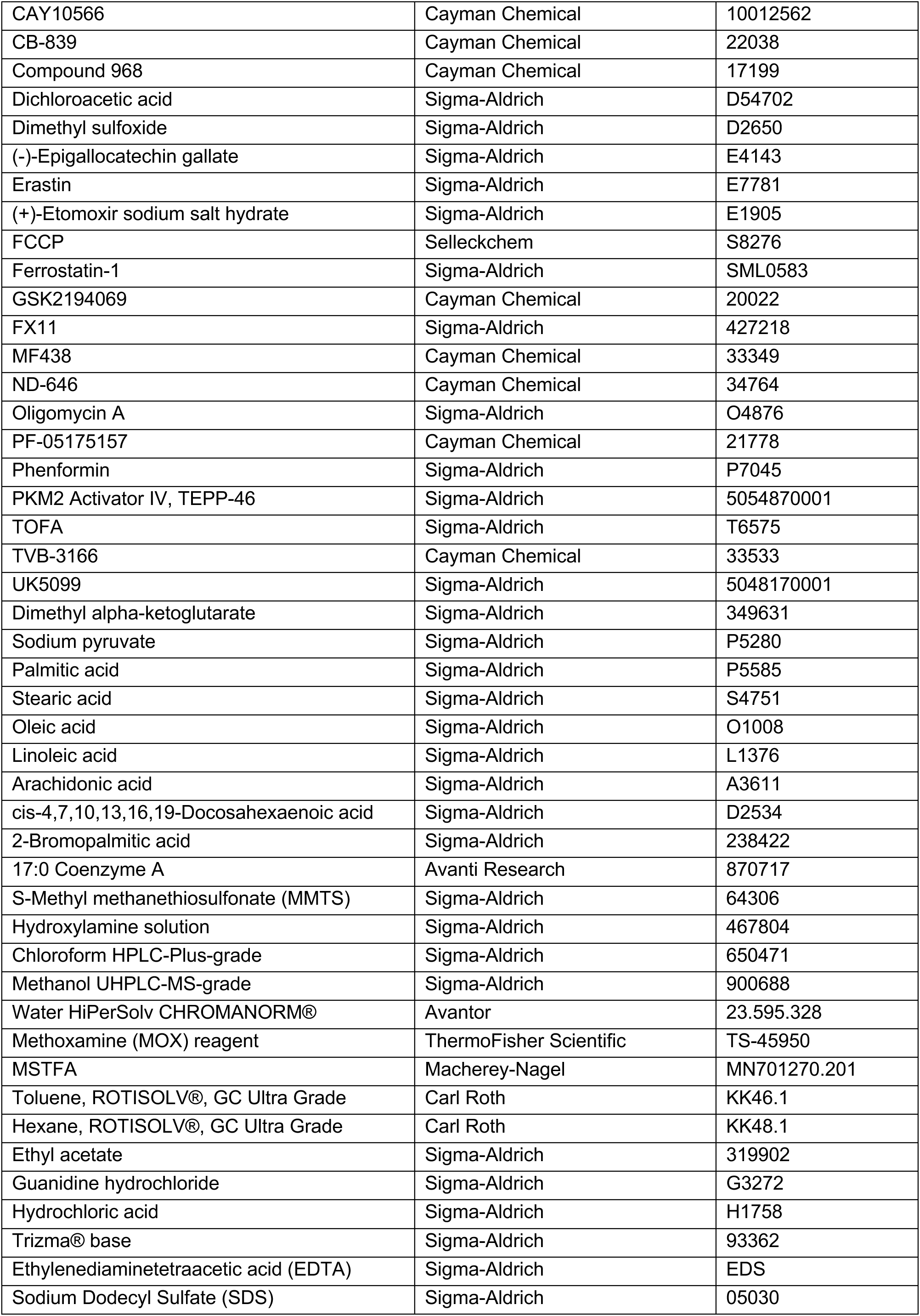
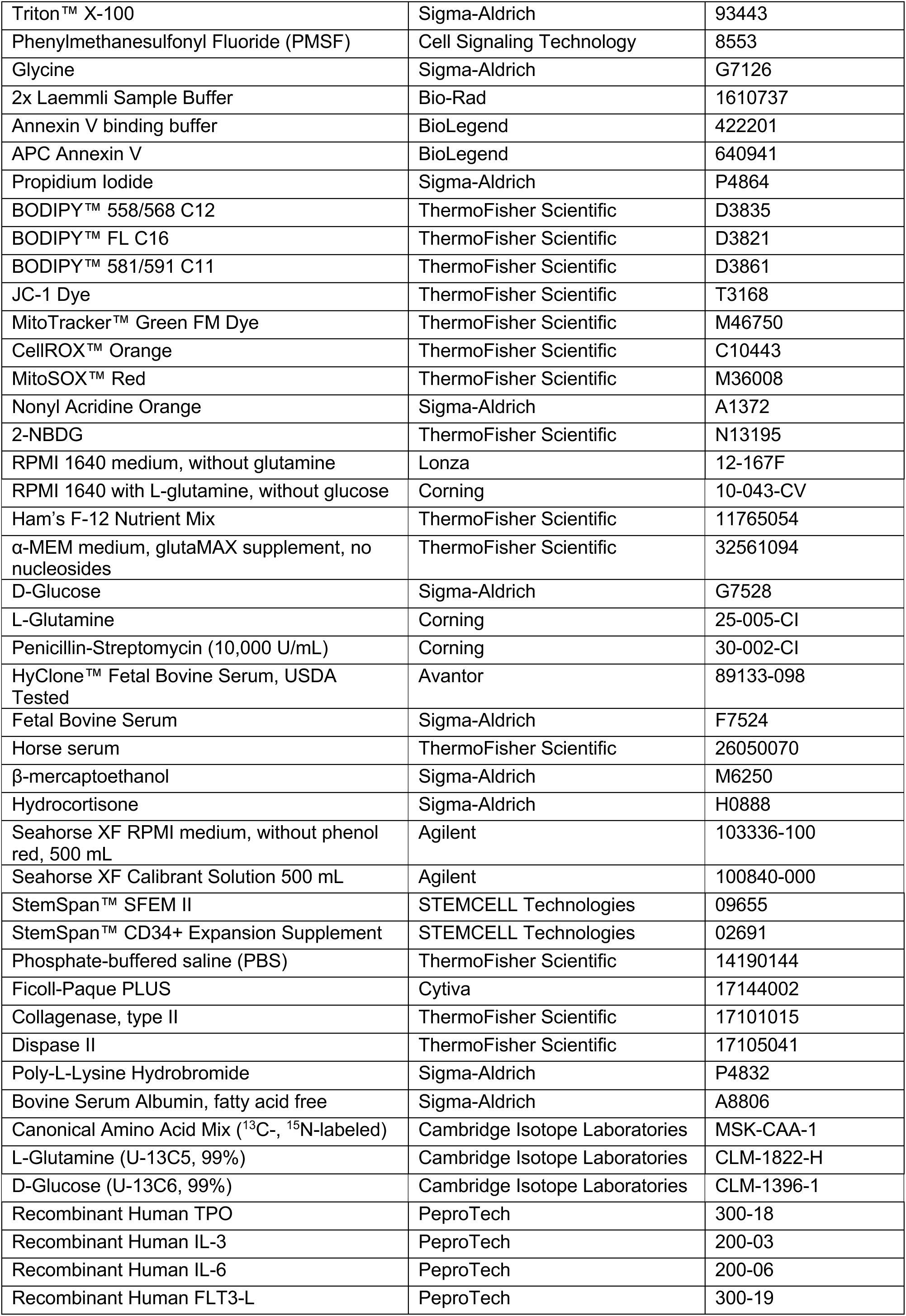
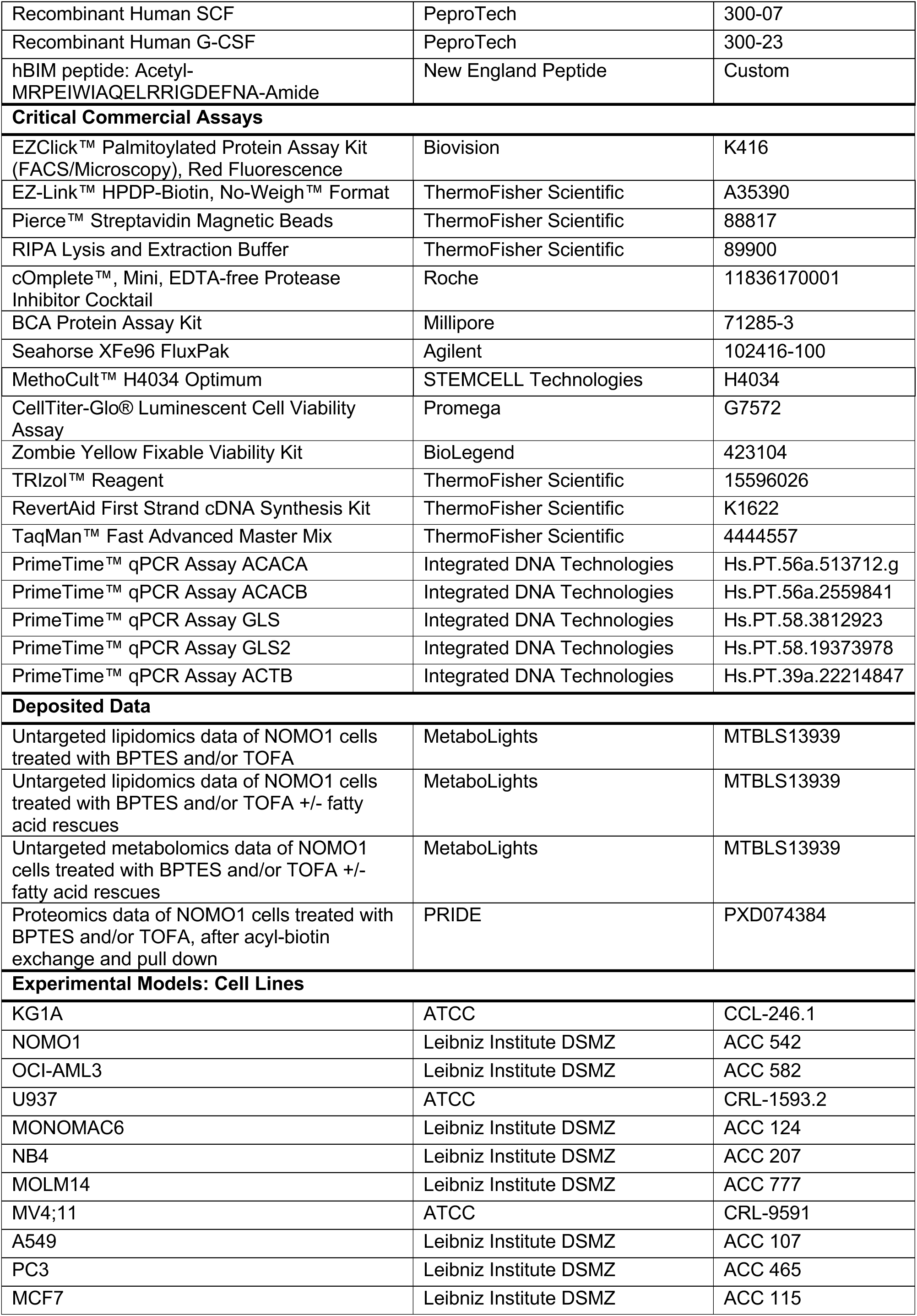
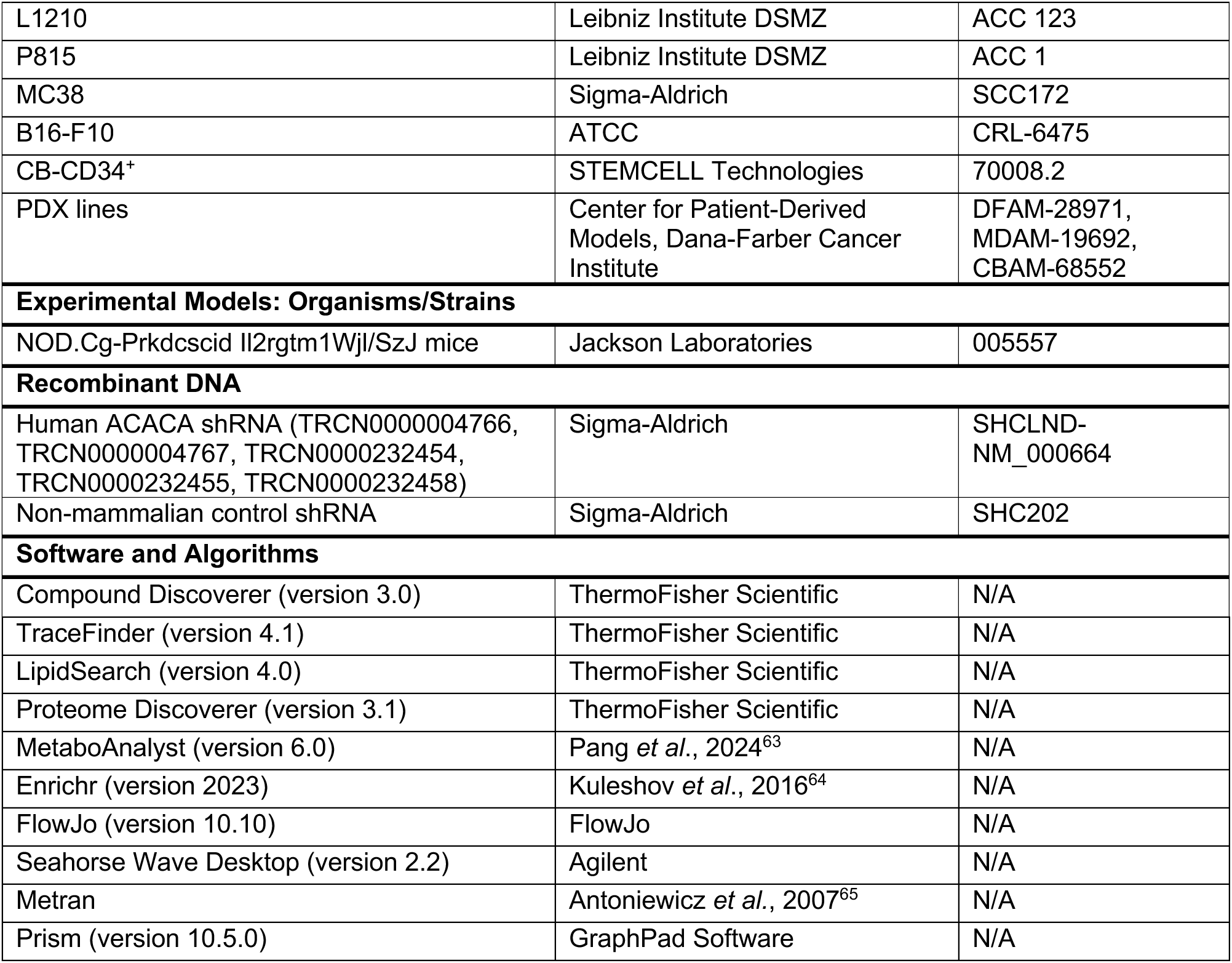

## RESOURCE AVAILABILITY

### Lead Contact

Further information and requests for resources and reagents should be directed to and will be fulfilled by the Lead Contact, Nick van Gastel.

### Materials Availability

This study did not generate new unique materials.

### Data and Code Availability

The mass spectrometry lipidomics and metabolomics data that support the findings of this study have been deposited in the MetaboLights^66^ database with the dataset identifier MTBLS13939 (https://www.ebi.ac.uk/metabolights/MTBLS13939). The mass spectrometry proteomics data have been deposited to the ProteomeXchange Consortium via the PRIDE^67^ partner repository with the dataset identifier PXD074384 and 10.6019/PXD074384. Genome-scale CRISPR screen data (DepMap Public 24Q2+Score, Chronos) and RNA-seq data (Batch-corrected TPM values) of AML cell lines (26 lines) were retrieved from the DepMap portal (https://depmap.org/portal)^31^. All other data supporting the findings of this study are available within the paper.

## EXPERIMENTAL MODEL AND SUBJECT DETAILS

### Mice

NOD.Cg-Prkdc^scid^ Il2rg^tm1Wjl^/SzJ (NSG) mice were purchased from Charles River. All animal procedures performed in this study were approved by the UCLouvain Institutional Animal Care and Use Committee.

### Patient Samples and Patient-Derived Xenografts

Healthy donor and AML patient bone marrow samples were collected with patient informed consent. Patient-derived xenograft (PDX) samples were obtained from the Center for Patient Derived Models at the Dana-Farber Cancer Institute and expanded in NSG mice. All studies were approved by the UCLouvain-Clinique Universitaire St. Luc, Massachusetts General Hospital or Dana-Farber Cancer Institute institutional review boards and conformed to guidelines for ethical research conduct in the Declaration of Helsinki.

### Cell Culture

All cell cultures were performed in an incubator at 37°C and 5% CO_2_. A549 cells were cultured in Ham’s F-12 Nutrient Mix supplemented with 2 mM L-glutamine, 100 U/mL penicillin, 100 µg/mL streptomycin and 10% heat-inactivated fetal bovine serum (FBS). All other cell lines were cultured in RPMI1640 medium supplemented with 2 mM L-glutamine, 100 U/mL penicillin, 100 µg/mL streptomycin and 10% heat-inactivated FBS. Cell line identity was validated by DNA fingerprinting and cultures were tested for mycoplasma contamination every six months using a Mycoplasma PCR Detection Kit (Abcam). CB-CD34^+^ cells were cultured in StemSpan SFEM II medium supplemented with 1X StemSpan CD34+ Expansion Supplement. For tracing experiments, CB-CD34^+^ cells were switched to RPMI160 medium supplemented with 2 mM L-glutamine, 100 U/mL penicillin, 100 µg/mL streptomycin, 10% heat-inactivated FBS, 50 ng/mL SCF, 50 ng/mL FLT3-L, 20 ng/mL TPO and 10 ng/mL IL-6 (50% of both media one week before and 100% RPMI1640 2 days before initiation of the tracing experiment). AML PDX cells and primary AML patient samples were cultured in α-MEM medium supplemented with 100 U/mL penicillin, 100 µg/mL streptomycin, 12.5% heat-inactivated FBS, 12.5% heat-inactivated horse serum, 57.2 µM β-mercaptoethanol, 1 µM hydrocortisone, 20 ng/mL G-CSF, 20 ng/mL TPO and 20 ng/mL IL-3.

Primary human BM-MNCs and BMSCs were isolated from bone marrow isolates obtained during routine hip arthroplasty. Samples were first passed through a 40 µm cell strainer. For the isolation of BM-MNCs, the flow-through was collected and overlayed onto Ficoll-Paque PLUS density gradient medium. The buffy coat was collected, and cells were expanded in RPMI160 medium supplemented with 2 mM L-glutamine, 100 U/mL penicillin, 100 µg/mL streptomycin, 10% heat-inactivated FBS, 50 ng/mL SCF, 50 ng/mL FLT3-L, 20 ng/mL TPO and 10 ng/mL IL-6. For BMSCs, the fraction retained on the cell strainer was digested with 3 mg/ml collagenase type II and 4 mg/ml dispase II in RPMI1640 medium for 30 minutes at 37°C in a shaking water bath, after which cells were passed through a 70 µm cell strainer and the flow-through was plated in α-MEM medium supplemented with 100 U/mL penicillin, 100 µg/mL streptomycin, 10% heat-inactivated FBS and 10% heat-inactivated horse serum.

Unless otherwise indicated, BPTES and TOFA were used at a final concentration of 5 µM in all experiments. DMSO was used as vehicle control. For fatty acid supplementation experiments, fatty acids (dissolved in ethanol) or vehicle (ethanol) were first complexed to fatty acid-free bovine serum albumin (4% in saline) for 1 hour at 37°C in a shaking water bath. Fatty acids were then added at 15 µM each to the culture medium.

## METHOD DETAILS

### Flow Cytometry

Cell viability was analyzed after staining of cells for 15 minutes with Annexin V-APC (1/20) and propidium iodide (PI; 1/2,000) or DAPI (1/20,000) in Annexin V binding buffer, with Annexin V^-^PI^-^ cells considered as viable cells. For the combinatorial metabolic compound screening cells were stained with PI directly added to the culture medium (1/2,000 final dilution) and PI^-^ cells were considered as viable cells. The number of viable cells per well was determined using Precision Count Beads™ (BioLegend). Apoptosis was measured using the PE Active Caspase-3 Apoptosis Kit (BD Biosciences) following the manufacturer’s instructions. The percentage of CD34^+^ was analyzed by staining cells with Alexa Fluor 488 anti-human CD34 (1/200) for 20 minutes in flow buffer (2% FBS in PBS). For lipid uptake, BODIPY™ 558/568 C12 or BODIPY™ FL C16 was added to the culture medium at 1 µM final concentration and incubated for 30 minutes, after which cells were washed with PBS and analyzed. To estimate glucose uptake, 2-NBDG was added to the culture medium at 100 ng/mL final concentration and incubated for 30 minutes, followed by washing and analysis. Mitochondrial parameters were measured by adding JC-1 (mitochondrial membrane potential; 5 µg/mL final concentration), MitoTracker™ Green FM (mitochondrial mass; 10 nM final concentration), Nonyl Acridine Orange (cardiolipin content; 15 ng/mL final concentration), CellROX™ Orange (total cellular ROS; 2.5 µM final concentration) or MitoSOX™ Red (mitochondrial superoxide levels; 2.5 µM final concentration) to the culture medium, incubating cells for 30 minutes in the incubator, followed by washing with PBS and analysis. To measure lipid peroxidation, BODIPY™ 581/591 C11 was added to the culture medium at 1 µM final concentration and incubated for 30 minutes, after which cells were washed with PBS and analyzed. Protein *S*-palmitoylation was measured using the EZClick™ Palmitoylated Protein Assay Kit following the manufacturer’s instructions. For all flow cytometry experiments, single color controls were used to set compensations, and fluorescence minus one controls were used to set gates. Flow cytometry data was collected on a BD LSR II or BD LSR Fortessa flow cytometer and analyzed using the FlowJo software.

### Dynamic BH3 Profiling

BH3 peptide plates (96-well) were generated by preparing dilutions of hBIM peptide (3 µM to 0.003 µM final concentration), DMSO (negative control) or alamethicin (250 nM final concentration, positive control) at 2X concentration in 15 µL BH3 profiling buffer (150 mM mannitol, 10 mM HEPES-KOH pH 7.5, 150 mM KCl, 0.02 mM EDTA, 0.02 mM EGTA, 0.1% BSA, 5 mM succinate) containing 0.002% digitonin. Dynamic BH3 profiling was carried out, as previously described in detail^68^, on NOMO1 cells, CB-CD34^+^ cells, primary AML samples or healthy donor BM-MNCs by treating cells for 18 hours with BPTES and/or TOFA at 5 µM each (or DMSO as vehicle). Next, cells were harvested and washed with PBS. For primary samples, cells were then resuspended in flow buffer (2% FBS in PBS) containing PE anti-human CD33, APC/Cy7 anti-human CD34, Alexa Fluor 488 anti-human CD38, Brilliant Violet 421 anti-human CD45 and Zombie Yellow Fixable Viability dye, each at 1:100 dilution, incubated for 30 minutes at 4°C and washed with PBS. Cells were then resuspended in BH3 profiling buffer, plated into a BH3 peptide plate at 5 × 10^4^ cells in 15 µL buffer per well, and incubated for 50 minutes in an incubator at 26°C. Next, cells were fixed by adding 10 µL of 4% paraformaldehyde to each well and incubated at room temperature for 10 minutes. Fixation was stopped by adding 10 µL of a 1.7 M Tris buffer containing 1.25 M Glycine (pH 9.1) to each well and samples were incubated at room temperature for 10 minutes. Next, 10 µL of pre-made Cytochrome C staining buffer (10% BSA, 2% Tween20 in PBS, with 1:400 dilution of anti-Cytochrome C-Alexa fluor 647 antibody) was added to the wells and samples were incubated overnight at 4°C in the dark. Data was collected on BD LSR Fortessa flow cytometer and analyzed using the FlowJo software. AML blasts were identified from viable cells as CD45^+^CD33^+^ and LSCs as CD45^+^CD34^+^CD38^-^ and the Mean Fluorescence Intensity (MFI) for Cytochrome C in each population was analyzed. Apoptotic priming was calculated for each concentration of hBIM peptide using the following formula:

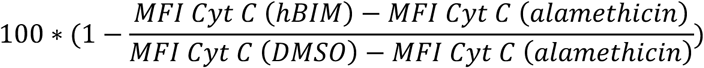

For each cell type/sample, the concentration of hBIM was then selected at which around 10% of Cytochrome C was released in vehicle-treated samples, and delta priming was calculated for BPTES-and/or TOFA-treated samples relative to vehicle-treated samples.

### Immunoblotting

Cells were lysed in RIPA buffer supplemented with 1x cOmplete protease inhibitor cocktail. Protein concentration was determined by bicinchoninic acid (BCA) assay, and equal amounts of proteins were separated by SDS-PAGE and transferred to a PVDF membrane with 0.45 µm pore size. Membranes were blocked with 5% dry milk in Tris-buffered saline (TBS) for 1 hour at room temperature and incubated overnight at 4°C with primary antibodies (all at 1/1,000 except for anti-β-actin which was used at 1/50,000) diluted in 5% BSA in TBS. Signals were detected using a LI-COR Odyssey CLx imaging system after incubation with IRDye-conjugated secondary antibodies.

### Quantitative RT-PCR

Total RNA was extracted using TRIzol reagent, and 1 μg of the extracted RNA was converted into cDNA using the RevertAid First Strand cDNA synthesis kit. The PCR reaction was performed using the TaqMan Fast Advanced Master Mix with pre-designed PrimeTime qPCR Assays with 6-FAM/ZEN/IBFQ reporter/quencher. Assays were performed in technical duplicate, and data were analyzed by the 2^-ΔCt^ method relative to *ACTB*.

### shRNAs

Cells were transduced with lentiviruses carrying shRNAs against human ACACA or a control shRNA (non-mammalian) through spinfection (1000 × g for 90 minutes at 30°C) in the presence of 8 µg/mL polybrene. After transduction, cells were selected with 5 µg/mL puromycin for at least 72 hours. Knockdown efficiency was verified through immunoblot (see Figure S4B).

### Stable Isotope Tracing by Gas Chromatography-Coupled Mass Spectrometry (GC-MS)

Cells were plated at 5 × 10^5^ cells in 1 mL RPMI1640 medium containing 2 mM U-^13^C_5_-glutamine or 11 mM U-^13^C_6_-glucose, as well as BPTES and/or TOFA at 5 µM each, or DMSO as vehicle. To correct for natural ^13^C abundances, control wells were included with cells cultured in medium supplemented with unlabeled glutamine and glucose. After 24 hours, cells were collected, washed with saline (0.9% NaCl in water), lysed in 400 µL UHPLC-grade methanol (-20°C) and transferred to a glass vial. To this, 200 µL ice-cold UHPLC-grade water and 400 µL UHPLC-grade chloroform were added, and samples were vortexed for 10 minutes at 4°C. Next, samples were centrifuged for 10 minutes at 5,000 × g, and the upper phase (containing polar metabolites) and lower phase (containing nonpolar metabolites) were collected separately and dried under nitrogen gas flow. Polar metabolites were resuspended in 15 µL of MOX reagent and incubated for 90 minutes at 30°C, with shaking at 300 rpm. Next, 30 µL of MSTFA was added, samples were vortexed and incubated for 30 minutes at 37°C. Samples were then centrifuged at 14,000 × g for 10 minutes and 40 µL were transferred into a GC-MS vial with glass insert. Nonpolar metabolites were resuspended in 100 µL toluene to which 750 µL methanol and 150 µL 8% hydrochloric acid in methanol were added. Samples were vortexed and incubated overnight at 45°C, after which 500 µL hexane and 500 µL water were added and samples were vortexed again. Samples were then allowed to sediment for 10 minutes at room temperature, after which the upper phase was transferred to a clean glass vial containing 500 µL water. This sedimentation step was repeated once more, after which the upper phase was transferred to a clean vial and dried under nitrogen gas flow. Nonpolar metabolites were then resuspended in 100 µL hexane and transferred into a GC-MS vial with glass insert. Samples were then analyzed on an Agilent 6890N GC equipped with DB-35ms capillary column (30 m × 0.25 mm × 0.25 μm) and 5975B Inert XL MS system under electron ionization at 70 eV and quadrupole mass analyzer. The MS source and quadrupole were held at 230°C and 150°C, respectively. Helium was used as carrier gas at a flow rate of 1 mL/minute. The mass isotopomer distribution was obtained by integrating the ion chromatograms over the full peak range for each set of metabolite mass fragments and was corrected for natural ^13^C abundance. To calculate the fraction of palmitate newly synthesized from glucose after tracer addition we performed isotopomer spectral analysis^69^ using the Metran software, as previously described^70^.

### Untargeted Lipidomics Analysis by Liquid Chromatography-Coupled MS (LC-MS)

After 24 hours of treatment with BPTES and or TOFA at 5 µM each, 10 million cells were collected per condition and washed with saline. Next, cells were lysed in 1 mL UHPLC-grade methanol (-20°C) and transferred to a 5 mL glass vial. To this, 500 µL ice-cold UHPLC-grade water and 2 mL UHPLC-grade chloroform were added, and samples were vortexed for 10 minutes at 4°C. Next, samples were centrifuged for 10 minutes at 5,000 × g, and the lower phase containing nonpolar metabolites was transferred to a clean glass vial and dried under nitrogen gas flow. Samples were then resuspended in 100 µL chloroform and split over two LC-MS vials for analysis in positive and negative mode separately. Samples were run on a ThermoFisher Ultimate 3000 UHPLC with Bio-Bond 5 µm C4 (50 x 4.6 mm) column (Dikma), coupled to a ThermoFisher Q Exactive plus Orbitrap MS with HESI source. For each sample 15 µL was injected, followed by gradient elution and MS acquisition. For positive mode, mobile phase A consisted of 5 mM ammonium formate, 0.1% formic acid and 5% methanol in water, and mobile phase B of 5 mM ammonium formate, 0.1% formic acid, 5% water, 35% methanol and 60% isopropanol. For negative mode, mobile phase A consisted of 0.03% ammonium hydroxide and 5% methanol in water, and mobile phase B of 0.03% ammonium hydroxide, 5% water, 35% methanol and 60% isopropanol. Gradients were as follows: minute 0-5, 0% B at 0.1 mL/min; minute 5-55, 20% to 100% B at 0.4 mL/min; minute 55-63, 100% B at 0.5 mL/min; minute 63-70: 0% B at 0.5 mL/min. MS parameters (Full MS to ddMS2): Full MS: 70,000 resolution, AGC target 1 × 10^6^, m/z range 150 to 2000; ddMS2: 35,000 resolution, AGC target 1 × 10^5^, loop count 5. Data was analyzed by the LipidSearch software for identification and integration of lipids, and datasets were further curated manually.

### Untargeted Metabolomics Analysis by LC-MS

After 24 hours of treatment with BPTES and or TOFA at 5 µM each, 5 million cells were collected per condition and washed with saline. Next, cells were lysed in 1 mL UHPLC-grade methanol (-20°C) and transferred to a 5 mL glass vial. To this, 500 µL ice-cold UHPLC-grade water and 1 mL UHPLC-grade chloroform were added, and samples were vortexed for 10 minutes at 4°C. Next, samples were centrifuged for 10 minutes at 5,000 × g, and the upper phase containing polar metabolites was transferred to a clean glass vial and dried under nitrogen gas flow. Samples were then resuspended in 100 µL of 70% acetonitrile in water containing 2.5 µM of an internal standard (^13^C-, ^15^N-labeled amino acid mix) and transferred to LC-MS vials. Samples were run on a ThermoFisher Ultimate 3000 UHPLC with ZIC-pHILIC 5 µm (150 x 2.1 mm) column (Merck), coupled to a ThermoFisher Q Exactive plus Orbitrap MS with HESI source. For each sample 5 µL was injected, followed by gradient elution and MS acquisition in polarity-switching mode. Mobile phase A consisted of 20 mM ammonium carbonate and 0.1% ammonium hydroxide in water, and mobile phase B of 97% acetonitrile in water. Gradients were as follows: minute 0-20, 100% to 40% B at 0.2 mL/min; minute 20-30, 40% to 0% B at 0.2 mL/min; minute 30-35, 0% B at 0.2 mL/min; minute 35-40: 0% to 100% B at 0.2 mL/min; minute 40-50: 100% B at 0.2 mL/min. MS parameters (Full MS): 70,000 resolution, AGC target 3 × 10^6^, m/z range 66.7 to 1000. A pooled sample was created from all samples and used for MS/MS runs. Data were processed with the Compound Discoverer software, including normalization by median centering to compensate for sample-to-sample variation and compound annotation using an in-house mzVault database and the mzCloud mass spectral library, and data visualization was achieved using the MetaboAnalyst software.

### Analysis of Protein-Bound Fatty Acids

After 24 hours of treatment with BPTES and or TOFA at 5 µM each, 5 million cells were collected per condition and washed three times with PBS and once with 1 M Tris buffer (pH 7.5). Fatty acids covalently bound to proteins were then isolated and analyzed following a previously published protocol^37^ with small adaptations. Cell pellets were resuspended in 500 µL SDS buffer (2% SDS, 62.5 mM Tris-HCl, pH 6.8) and heated at 95°C for 5 minutes while shaking (800 rpm). Samples were then transferred to 11 mL glass vials and 10 mL CHCl_3_/MeOH (2:1, vol/vol) was added. Samples were vortexed, incubated at room temperature for 30 minutes, and centrifuged at 4,815 × g at room temperature for 15 minutes. The supernatant was discarded and the extraction sequence was repeated on the pellet 4 more times with respectively 10 mL CHCl_3_/MeOH (2:1), CHCl_3_/MeOH (1:2), CHCl_3_/MeOH/H_2_O (1:1:0.3) and MeOH. The final precipitate was resuspended in 500 µL guanidine HCl (6 M) containing 20 mM Tris, 0.02% EDTA and 4 nmol C17:0-CoA (internal standard), and a 20 µL aliquot was taken and used for protein determination. To release the thioester-bound fatty acids from the proteins, 500 µL of hydroxylamine (50% in H_2_O) was added to the samples. A blank was created containing cells but without hydroxylamine, receiving 500 µL of 1 M Tris (pH 7.5) instead. Samples were then briefly vortexed and incubated at room temperature for 16 hours while shaking (500 rpm). Next, 1 mL CHCl_3_/ethyl acetate (1:1) was added, samples were vortexed for 1 minute and centrifuged at 4,815 × g at room temperature for 15 minutes, creating two phases. The infranatant was collected and transferred to a clean 5 mL glass vial, and 1 mL CHCl_3_/ethyl acetate (1:1) was added to the original tube, which was vortexed and centrifuged again. The resulting infranatant was added to the 5 mL glass vial containing the infranatant from the first extraction, and the whole procedure was repeated a third time. The three pooled infranatants were then centrifuged at 4,815 × g at room temperature for 15 minutes, and the infranatant was again collected, transferred to a clean 5 mL glass tube and dried completely under nitrogen flow. The samples were then resuspended in 200 µL CHCl_3_/ethyl acetate (1:1), vortexed and centrifuged at 1,115 × g at room temperature for 5 minutes. The supernatant was transferred to a clean tube, and the original tube was washed with an additional 200 µL CHCl_3_/ethyl acetate (1:1) which, after vortexing and centrifugation, was added to the first supernatant. The pooled supernatants were dried again under nitrogen flow, and the samples were resuspended in 100 µL toluene to which 750 µL methanol and 150 µL 8% hydrochloric acid in methanol were added. Samples were vortexed and incubated overnight at 45°C, after which 500 µL hexane and 500 µL water were added and samples were vortexed again. Samples were then allowed to sediment for 10 minutes at room temperature, after which the upper phase was transferred to a clean glass vial containing 500 µL water. This sedimentation step was repeated once more, after which the upper phase was transferred to a clean vial and dried under nitrogen gas flow. Derivatized fatty acids were then resuspended in 100 µL hexane and transferred into a GC-MS vial with glass insert. Samples were then analyzed on an Agilent 7890B GC equipped with CP-Sil 8 CB-MS capillary column (30 m × 0.25 mm × 0.25 μm) and 5977A MSD system. Ion chromatograms were integrated over the full peak range, blank values were subtracted, and values were corrected for sample protein content.

### Acyl-Biotin Exchange

The protocol for acyl-biotin exchange was adapted from Wan *et al*.^71^ and Tewari *et al*.^72^, and a detailed schematic overview of the procedure can be found in Figure S6A. After 24 hours of treatment, 20 million cells were collected per condition, washed with PBS and resuspended in 1 mL lysis buffer (150 mM NaCl, 50 mM Tris, 5 mM EDTA, pH 7.4) supplemented with 2 mM PMSF, 1.7% Triton X-100 and 1x cOmplete protease inhibitor cocktail, and samples were incubated at 4°C for 2 hours with rotation. After incubation, the lysates were pre-cleared by centrifuging at 12,000 × g for 5 minutes. Protein from the solution was precipitated with methanol (4 volumes), chloroform (1.5 volumes) and water (3 volumes), followed by centrifugation at 4,800 × g for 20 min. The interphase was collected, and three volumes of methanol were added, followed by centrifugation at 4,000 × g for 15 minutes. The pellet was air-dried, then dissolved in 200 μL of 2% SDS buffer (2% SDS, 50 mM Tris, 5 mM EDTA, pH 7.4). Free thiols were blocked by adding 200 µL of 2% SDS buffer containing 2% MMTS, and samples were incubated at 42 °C for 15 minutes in a thermal shaker. To remove unbound MMTS, the samples were subjected to three chloroform-methanol precipitation washes as described above. After the third and last wash, the pellet was air-dried and dissolved in 200 µL 4% SDS buffer (4% SDS, 50 mM Tris, 5 mM EDTA, pH 7.4). Hydroxylamine cleavage and capture of *S*-acylated proteins was performed by adding 1 mL of HA buffer (0.7 M hydroxylamine, 1 mM HPDP-biotin, pH 7.4, supplemented with 0.2% Triton X-100, 1 mM PMSF and 1x cOmplete protease inhibitor cocktail) and incubating at room temperature for 2 hours with rotation. A negative control was created containing untreated cells but without hydroxylamine (receiving 1 mL of 50 mM Tris, 1 mM HPDP-biotin, pH 7.4, supplemented with 0.2% Triton X-100, 1 mM PMSF and 1x cOmplete protease inhibitor cocktail) and a blank was created containing untreated cells but without HPDP-biotin (receiving 1 mL of 0.7 M hydroxylamine, 50 mM Tris, pH 7.4, supplemented with 0.2% Triton X-100, 1 mM PMSF and 1x cOmplete protease inhibitor cocktail. To remove unbound HPDP-biotin, three chloroform-methanol washes were performed, and the pellet was dissolved in 120 μL of 2% SDS buffer. Protein concentration was determined by BCA assay, and 1 mg of protein was incubated with 100 µL streptavidin magnetic beads (10 mg/mL) in lysis buffer supplemented with 0.1% SDS and 0.1% Triton X-100 at room temperature for 2 hours with rotation. Unbound proteins next were removed using a magnetic stand, and the samples were washed five times using lysis buffer containing 0.1% SDS and 0.1% Triton X-100. For MS, the proteins were eluted in 100 µL of 0.2 M glycine. For immunoblots, the proteins were eluted in 100 μL of 2x Laemmli sample buffer containing 1% β-mercaptoethanol.

### Proteomics Analysis

The eluted proteins were precipitated by adding 15% trichloroacetic acid, samples were dried in a SpeedVac vacuum concentrator and resuspended in 10 µL of 50 mM ammonium bicarbonate, pH 8.0 with 0.5 µg trypsin (Promega). After overnight digestion at 37°C, the digestion was stopped by addition of trifluoroacetic acid (0.1% final). Peptides were dissolved in solvent A (0.1% trifluoroacetic acid in 2% acetonitrile) and directly loaded onto a reversed-phase pre-column (Acclaim PepMap 100, ThermoFisher Scientific) and eluted in backflush mode. Peptide separation was performed using a reversed-phase analytical column (EasySpray PepMap RSLC, 0.075 x 250 mm, ThermoFisher Scientific) with a linear gradient of 4%-27.5% solvent B (0.1% formic acid in 80% acetonitrile) for 40 minutes, 27.5%-50% solvent B for 20 minutes, 50%-95% solvent B for 10 minutes and holding at 95% for the last 10 minutes at a constant flow rate of 300 nL/min on a ThermoFisher Vanquish Neo UHPLC system coupled to a ThermoFisher Orbitrap Fusion Lumos or Orbitrap Exploris 240 MS with NSI source. Intact peptides were detected in the Orbitrap at a resolution of 120,000 (Lumos) or 60,000 (Exploris). Peptides were selected for MS/MS using HCD setting at 30, ion fragments were detected in the IonTrap (Lumos) or in the Orbitrap (Exploris) at a resolution of 30,000. A data-dependent procedure that alternated between one MS scan followed by MS/MS scans was applied for 3 seconds for ions above a threshold ion count of 1.0 × 10^4^ in the MS survey scan with 30-second dynamic exclusion. MS1 spectra were obtained with an AGC target of 4 × 10^5^ ions and a maximum injection time set to auto, and MS2 spectra were acquired with an AGC target of 1 × 10^4^ (IonTrap) or 5 × 10^4^ (Orbitrap) ions and a maximum injection set to auto. For MS scans, the m/z scan range was 350 to 1800. The resulting MS/MS data was processed using Sequest HT search engine within Proteome Discoverer against a *Homo sapiens* protein database obtained from Uniprot. Trypsin was specified as a cleavage enzyme allowing up to 2 missed cleavages, 3 modifications per peptide and up to 5 charges. Mass error was set to 10 ppm for precursor ions and 0.2 Da (IonTrap) or 10 ppm (Orbitrap) for fragment ions. Oxidation on Met (+15.995 Da), conversion of Gln (-17.027 Da) or Glu (- 18.011 Da) to pyro-Glu at the peptide N-term were considered as variable modifications. False discovery rate (FDR) was assessed using Percolator and thresholds for protein, peptide and modification site were specified at 1%. Label-free quantification (AUC) was performed in Proteome Discoverer after normalization on the total peptide amount. Data were further normalized by Centered Log-Ratio (CLR) transformation. Enrichment analysis of proteins included in different classes was performed in Enrichr^64^.

### Extracellular Flux Analysis

The oxygen consumption rate (OCR) and extracellular acidification rate (ECAR) were measured using a Seahorse XF96 analyzer (Seahorse Bioscience, Agilent) at 37 °C. For AML cell lines 1 × 10^5^ cells and for PDX samples 2 × 10^5^ cells were seeded per well in poly-L-lysine-coated Seahorse XF96 plates in 180 µL XF RPMI medium. For adherent tumor lines, 1 × 10^5^ cells were seeded in uncoated Seahorse XF96 plates in regular culture medium and incubated for 24 hours, after which they were washed with PBS and 180 µL XF RPMI medium was added to each well. For the MitoStress, ATP Rate and FuelFlex assays, XF RPMI medium was supplemented with 10 mM D-glucose and 2 mM L-glutamine. For the GlycoStress assay, XF RPMI medium was supplemented with 2 mM L-glutamine. Plates were incubated for 60 minutes at 37°C in a CO_2_-free incubator to stabilize the pH. Analyses were performed under basal conditions and after the following sequential injections: MitoStress assay AML cell lines: 1 µM oligomycin A (Port A), 1 µM FCCP (Port B) and 0.5 µM antimycin A (Port C); MitoStress assay solid tumor cell lines and PDX lines: 2.5 µM oligomycin A (Port A), 2.5 µM FCCP (Port B) and 2 µM antimycin A + 2 µM rotenone (Port C); GlycoStress assay: 10 mM glucose (Port A), 2.5 µM oligomycin A (Port B) and 50 mM 2-deoxy-D-glucose (Port C); ATP Rate assay: 1.5 µM oligomycin A (Port A), and 0.5 µM antimycin A + 0.5 µM rotenone (Port B); FuelFlex assay: etomoxir (50 µM), BPTES (5 µM) and UK5099 (2 µM) were sequentially injected alone or in combination using Ports A and B according to the manufacturer’s protocol, followed by 0.5 µM antimycin A (Port C). The Seahorse Wave Desktop software was used to design assays and for data analysis, and measurements were normalized to the number of cells per well.

### Cellular ATP Content

After treatment, cells were harvested and counted using a Countess II system (ThermoFisher Scientific). Equal number of cells were subsequently used for determining cellular ATP levels using the CellTiter-Glo Luminescent Cell Viability Assay, according to the manufacturer’s instructions.

## QUANTIFICATION AND STATISTICAL ANALYSIS

All numerical results are reported as mean ± standard deviation (SD). Statistical significance of the difference between experimental groups was analyzed by two-tailed unpaired Student’s t-test or by one-way or two-way ANOVA with Tukey or Dunnett’s multiple comparison test using the GraphPad PRISM 5 software. Differences were considered statistically significant for P < 0.05.

The coefficient of drug interaction (CDI) was calculated as follows:

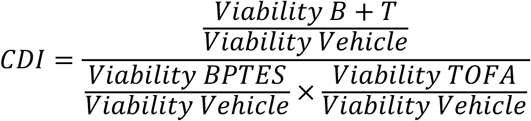

The following criteria were used to determine the type of drug interaction: CDI < 1: Synergism (significant if < 0.7); CDI = 1: Additive effect; CDI > 1: Antagonism.

## Supplemental Tables

Table S1. Genetic profile of AML patient samples and PDXs. Related to Figures 1, 6, S6.

Table S2. Lipidomics NOMO1 BPTES-TOFA. Related to Figures 3, S3.

Table S3. Lipidomics NOMO1 BPTES-TOFA FA rescues. Related to Figures 3, S3.

Table S4. Enzymes using FA-CoA. Related to Figure 5.

Table S5. ABE Proteomics NOMO1 BPTES-TOFA. Related to Figures 6, S6.

Table S6. ANOVA Top100 ABE Proteomics NOMO1 BPTES-TOFA. Related to Figures 6, S6.

Table S7. Metabolomics NOMO1 BPTES-TOFA FA rescues. Related to Figure 6.

## Supplemental Figures

**Figure S1.**
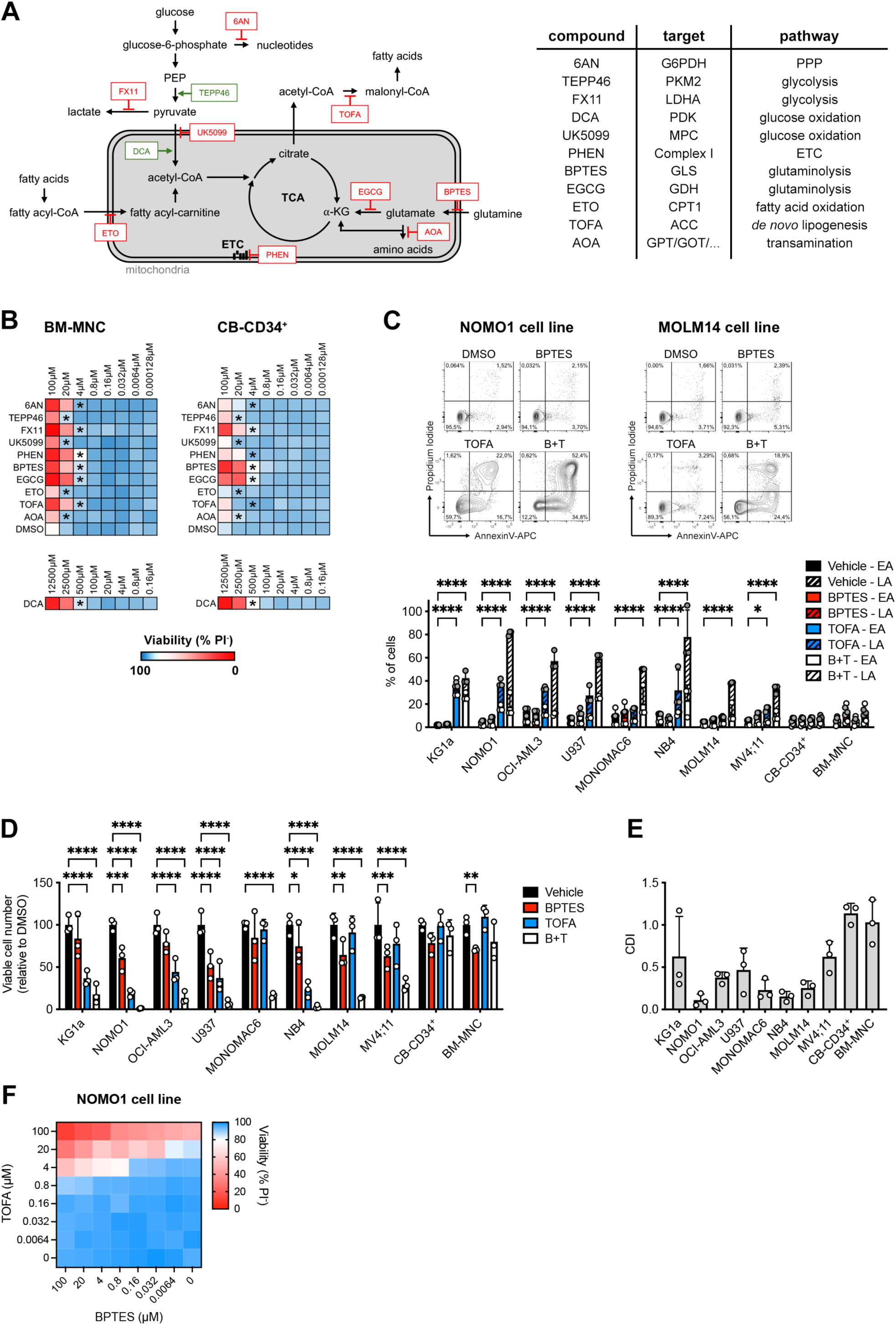
BPTES and TOFA Induce Apoptosis in AML Cells (related to Figure 1). (**A**) Schematic showing the expected metabolic targets of the inhibitors used in the combinatorial screen. (**B**) Heatmaps showing viability of bone marrow mononuclear cells (BM-MNC) and cord blood-derived CD34^+^ cells (CB-CD34^+^) treated for 72 hours with increasing doses of different metabolic inhibitors. Stars indicate the concentrations where both healthy hematopoietic stem/progenitor cell types remained above 80% viable, and which were used for the combinatorial compound screening on AML cell lines. (**C**) Apoptosis of AML cell lines and healthy hematopoietic stem/progenitor cells treated for 72 hours with BPTES (5 µM) and/or TOFA (5 µM). Plots on top show representative samples from NOMO1 and MOLM14 cell lines. EA, early apoptotic (AnxV^+^PI^-^); LA, late apoptotic (AnxV^+^PI^+^). (**D**) Viable cell number in cultures of AML cell lines and healthy hematopoietic stem/progenitor cells treated for 72 hours with BPTES and/or TOFA. (**E**) Coefficient of Drug Interaction (CDI) for BPTES and TOFA calculated from the viable cell count data shown in D. (**F**) Heatmap showing viability of NOMO1 AML cells treated for 72 hours with increasing doses of BPTES and/or TOFA. *p < 0.05, **p < 0.01, ***p < 0.001, ****p < 0.0001.

**Figure S2.**
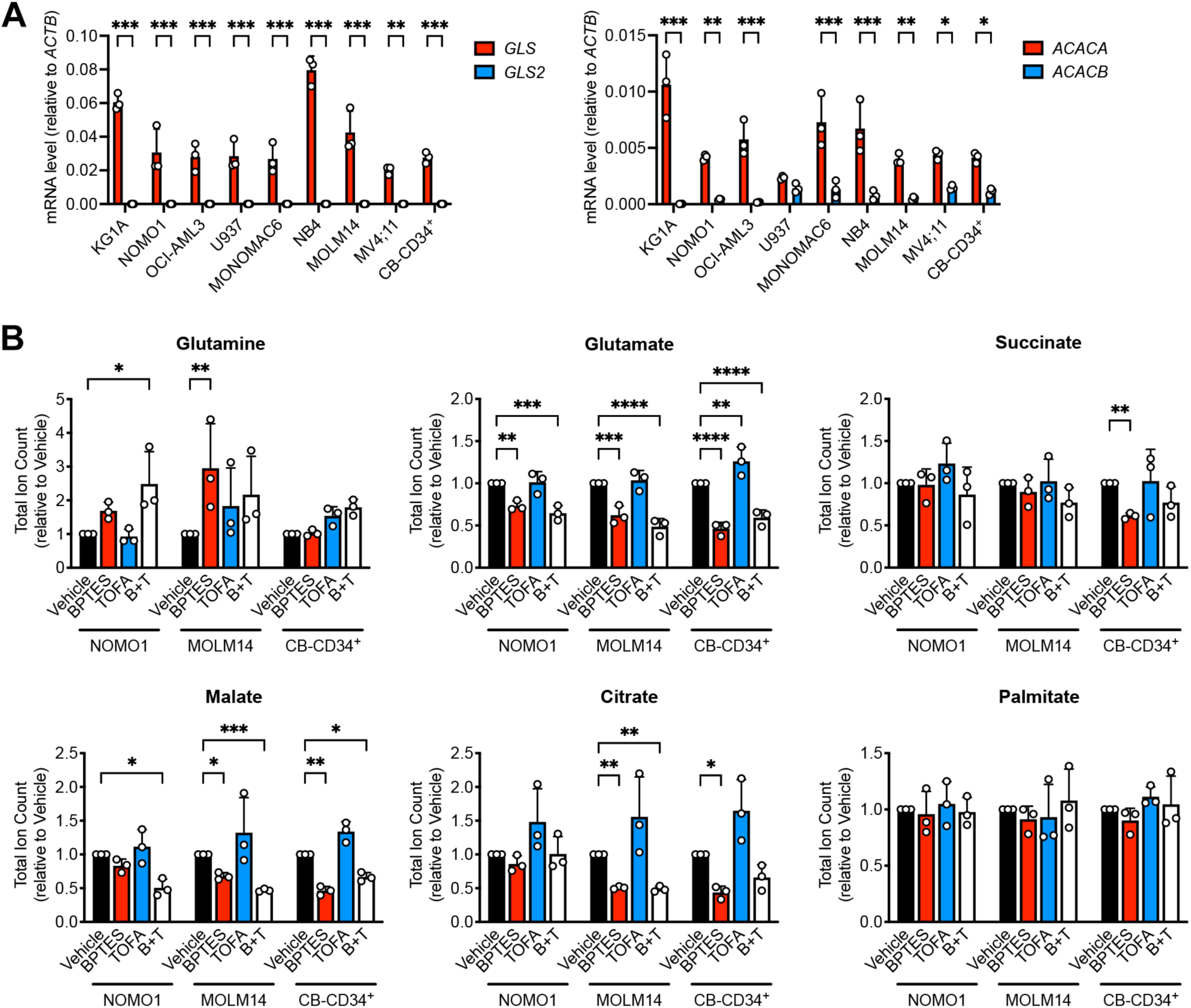
Metabolic Effects of BPTES and TOFA on AML Cells and HSPCs (related to Figure 2). (**A**) Gene expression of GLS isoforms (left) and ACC isoforms (right) in AML cell lines and cord blood-derived CD34^+^ cells (CB-CD34^+^). (**B**) Intracellular levels of glutamine, glutamate, succinate, malate, citrate and lipid palmitate in AML cell lines or cord blood-derived CD34^+^ cells (CB-CD34^+^) treated for 24 hours with BPTES (5 µM) and/or TOFA (5 µM). *p < 0.05, **p < 0.01, ***p < 0.001, ****p < 0.0001.

**Figure S3.**
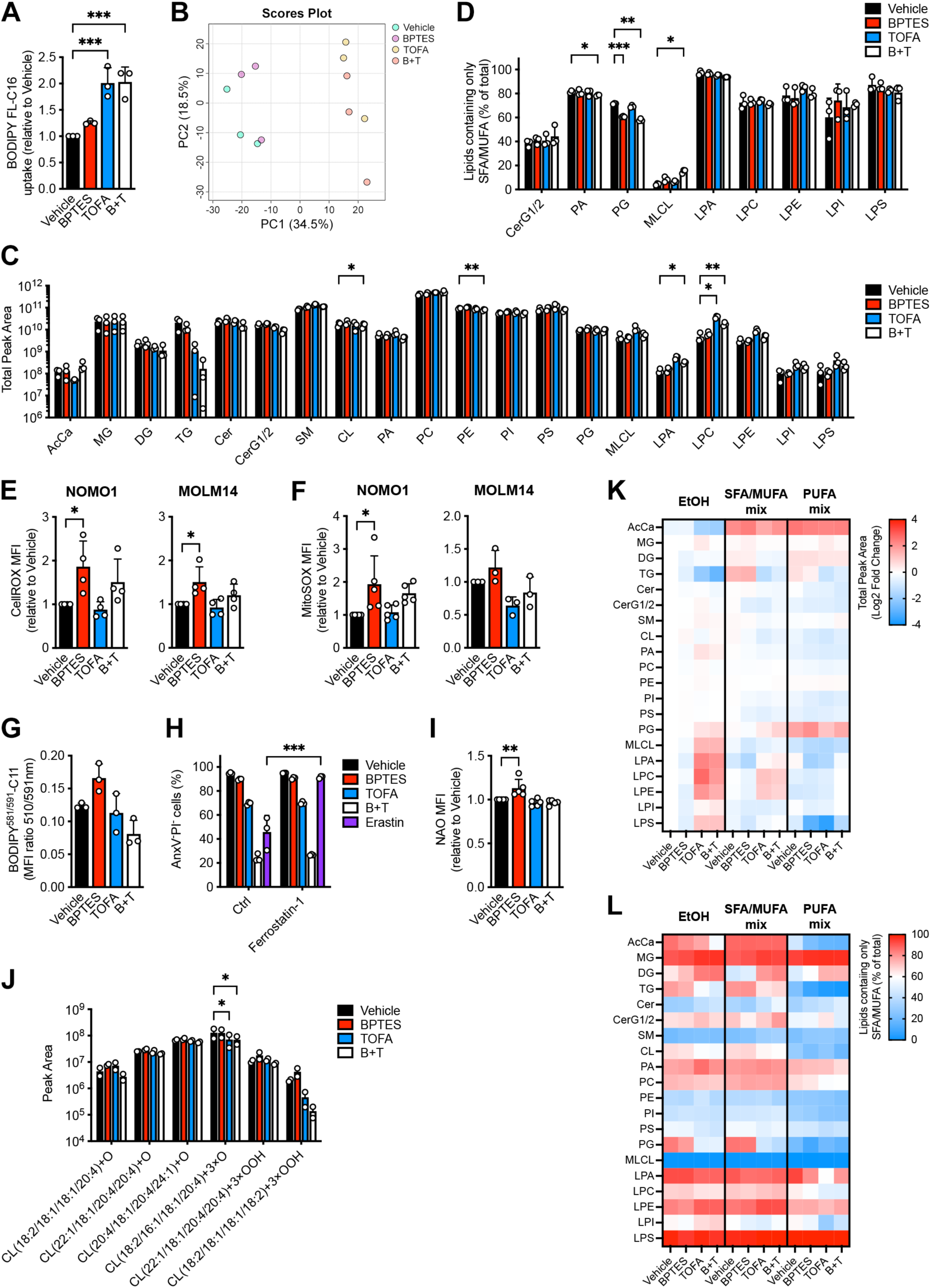
Lipid Profiling in AML Cells After BPTES and TOFA Treatment (related to Figure 3). (**A**) Uptake of a fluorescently labeled C16 fatty acid in NOMO1 AML cells treated for 24 hours with BPTES (5 µM) and/or TOFA (5 µM), relative to vehicle-treated cells. (**B**) Principal component analysis of lipidomics data of NOMO1 AML cells treated for 24 hours with BPTES and/or TOFA. (**C**) Lipid abundance across different lipid classes in NOMO1 AML cells treated for 24 hours with BPTES and/or TOFA, as determined by lipidomics. AcCa, acylcarnitines; MG, monoacylglycerides; DG, diacylglycerides; TG, triacylglycerides; Cer, ceramides; CerG1/2, mono/diglucosylceramides; SM, sphingomyelins; CL, cardiolipins; PA, phosphatidic acids; PC, phosphatidylcholines; PE, phosphatidylethanolamines; PI, phosphatidylinositols; PS, phosphatidylserines; PG, phosphatidylglycerols; MLCL, monolysocardiolipins; LPA, lysophosphatidic acids; LPC, lysophosphatidylcholines; LPE, lysophosphatidylethanolamines; LPI, lysophosphatidylinositols; LPS, lysophosphatidylserines. (**D**) Saturation degree across different lipid classes in NOMO1 AML cells treated for 24 hours with BPTES and/or TOFA, as determined by lipidomics. (**E-F**) Total cellular reactive oxygen species levels measured by CellROX staining (E), and mitochondrial superoxide levels measured by MitoSOX (F) in AML cell lines treated for 24 hours with BPTES and/or TOFA, relative to vehicle-treated cells. (**G**) Lipid peroxidation measured by BODIPY^581/591^-C11 staining in NOMO1 AML cells treated for 24 hours with BPTES and/or TOFA. MFI, mean fluorescence intensity. (**H**) Viability of NOMO1 AML cells treated for 72 hours with BPTES and/or TOFA, or with Erastin (1 µM), in the presence or absence of Ferrostatin-1 (1 µM). (**I**) Levels of non-oxidized cardiolipins measured by 10-N-nonyl acridine orange (NAO) staining in NOMO1 AML cells treated for 24 hours with BPTES and/or TOFA, relative to vehicle-treated cells. (**J**) Abundance of oxidized and peroxidized cardiolipins in NOMO1 AML cells treated for 24 hours with BPTES and/or TOFA, as determined by lipidomics. (**K-L**) Heatmaps showing lipid abundance (K) and saturation degree (L) across different (phospho)lipid classes in NOMO1 AML cells treated for 24 hours with BPTES and/or TOFA in the presence of exogenously added mixtures of albumin-complexed saturated/monounsaturated fatty acids (SFA/MUFA; palmitic acid, stearic acid and oleic acid at 15 µM each) or polyunsaturated fatty acids (PUFA; linoleic, arachidonic and docosahexaenoic acid at 15 µM each). Ethanol/albumin (EtOH) was used as vehicle. *p < 0.05, **p < 0.01, ***p < 0.001.

**Figure S4.**
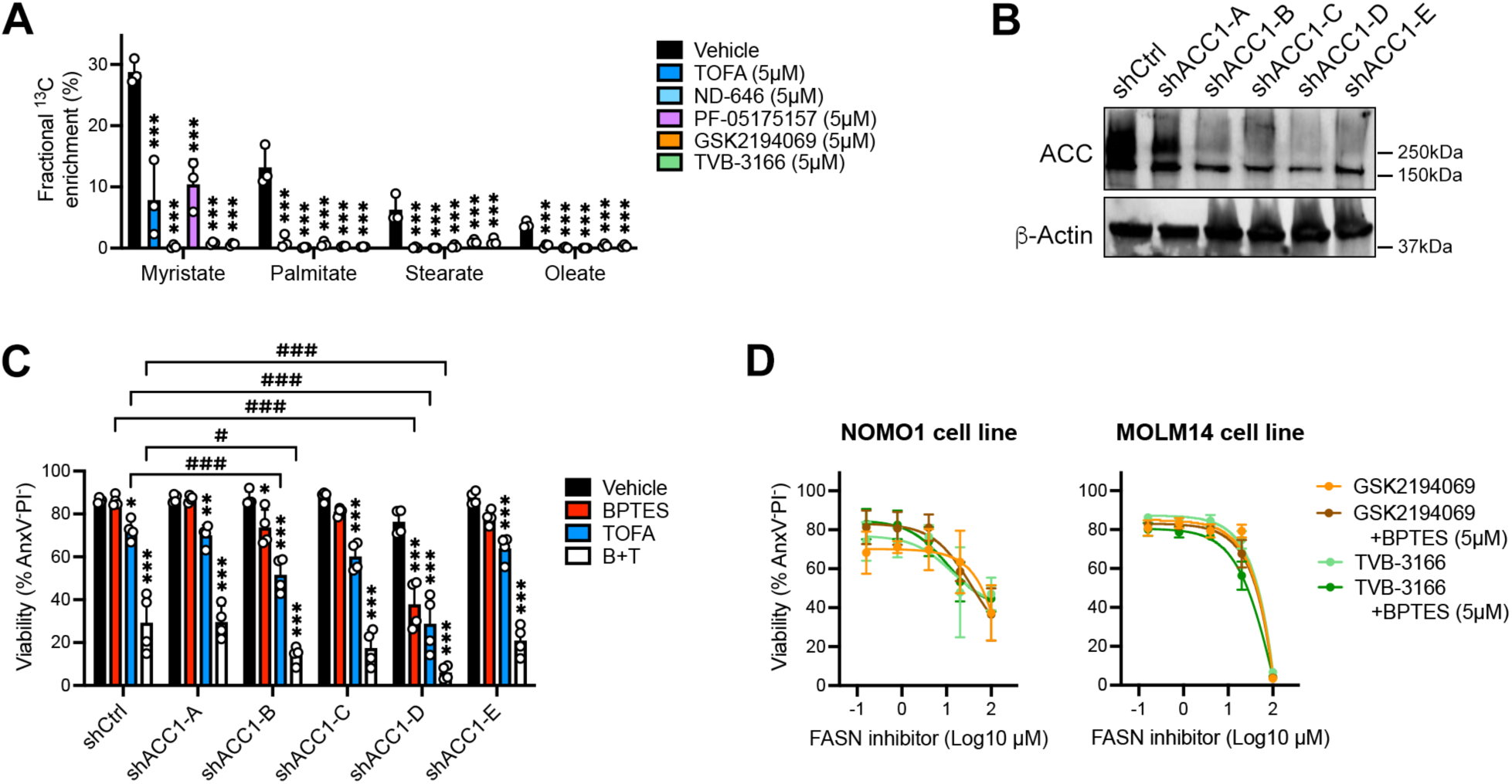
Inhibition of *De Novo* Lipogenesis in AML (related to Figure 4). (**A**) Total fractional enrichment from ^13^C_5_-glucose in intracellular lipid myristate, palmitate, stearate and oleate in NOMO1 AML cells treated for 24 hours with different inhibitors of *de novo* lipogenesis. (**B**) Immunoblot detection of total acetyl-CoA carboxylase (ACC) protein levels in NOMO1 AML cells transduced with lentiviruses carrying short hairpin RNAs (shRNAs) against human ACACA or a control shRNA (non-mammalian), with β-actin as loading control. (**C**) Viability of NOMO1 AML cells, transduced with lentiviruses carrying shRNAs against human ACACA or a control shRNA, treated for 72 hours with BPTES (5 µM) and/or TOFA (5 µM). (**D**) Viability of AML cell lines (mean ± SD; N=3) treated for 72 hours with increasing doses of GSK2194069 or TVB-3166 in the presence or absence of BPTES (5 µM). FASN, fatty acid synthase. *p<0.05, **p < 0.01, ***p < 0.001.

**Figure S5.**
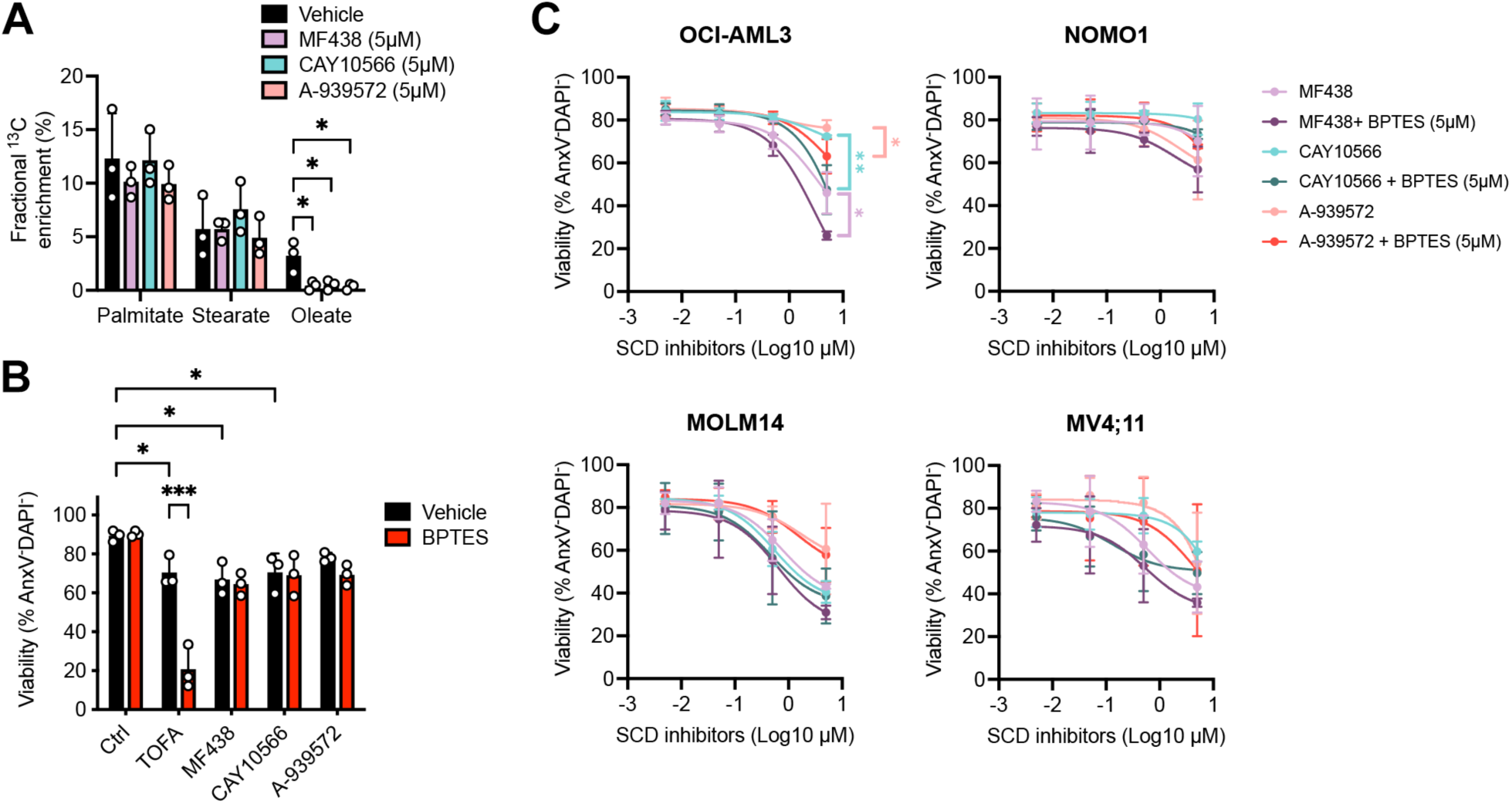
SCD Inhibition Does Not Sensitize AML Cells to BPTES (related to Figure 5). (**A**) Total fractional enrichment from ^13^C_5_-glucose in intracellular lipid palmitate, stearate and oleate in NOMO1 AML cells treated for 24 hours with different SCD inhibitors. SCD, stearoyl-CoA desaturase (**B**) Viability of NOMO1 AML cells treated for 72 hours with BPTES (5 µM) alone or in combination with TOFA (5 µM) or with different SCD inhibitors (5 µM). (**C**) Viability of AML cell lines (mean ± SD; N=3) treated for 72 hours with increasing doses of MF438, CAY10566 or A-939572 in the presence or absence of BPTES (5 µM). *p<0.05, **p < 0.01, ***p < 0.001.

**Figure S6.**
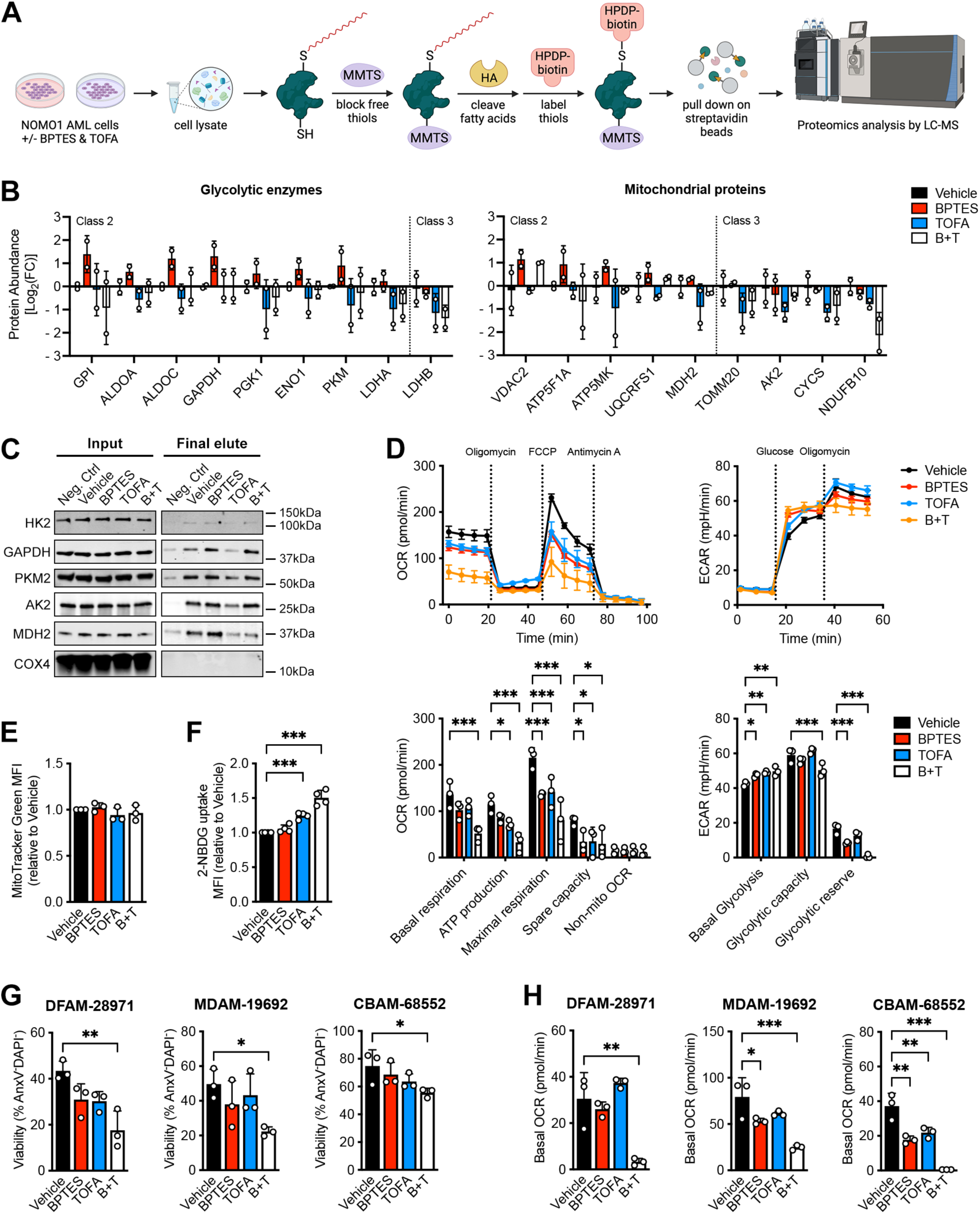
TOFA Prevents Metabolic Adaptation to Inhibition of Glutaminolysis (related to Figure 6). (**A**) Schematic showing the acyl-biotin exchange (ABE) assay procedure. (**B**) Changes in *S*-acylation of glycolytic enzymes (left) and mitochondrial proteins (right) in NOMO1 AML cells treated for 24 hours with BPTES (5 µM) and/or TOFA (5 µM), relative to vehicle-treated cells. (**C**) Immunoblot detection of total levels or *S*-acylated forms (obtained after ABE) of different metabolic enzymes in NOMO1 AML cells treated for 24 hours with BPTES and/or TOFA. (**D**) Real-time oxygen consumption rate (OCR; top left) and extracellular acidification rate (ECAR; top right) of MOLM14 AML cells (mean ± SD; N=3) treated for 24 hours with BPTES and/or TOFA, measured by Seahorse assay. Graphs show quantification of different OXPHOS parameters from OCR data (bottom left) and glycolysis parameters from ECAR data (bottom right). (**E-F**) Mitochondrial mass measured by MitoTracker Green staining (E), and uptake of a fluorescently labeled glucose analog (F) in NOMO1 AML cells treated for 24 hours with BPTES and/or TOFA. MFI, mean fluorescence intensity. (**G**) Viability of three different AML PDX lines treated for 72 hours with BPTES and/or TOFA. (**H**) Basal OCR of three different AML PDX lines treated for 24 hours with BPTES and/or TOFA, measured by Seahorse assay. *p < 0.05, **p < 0.01, ***p < 0.001.

**Figure S7.**
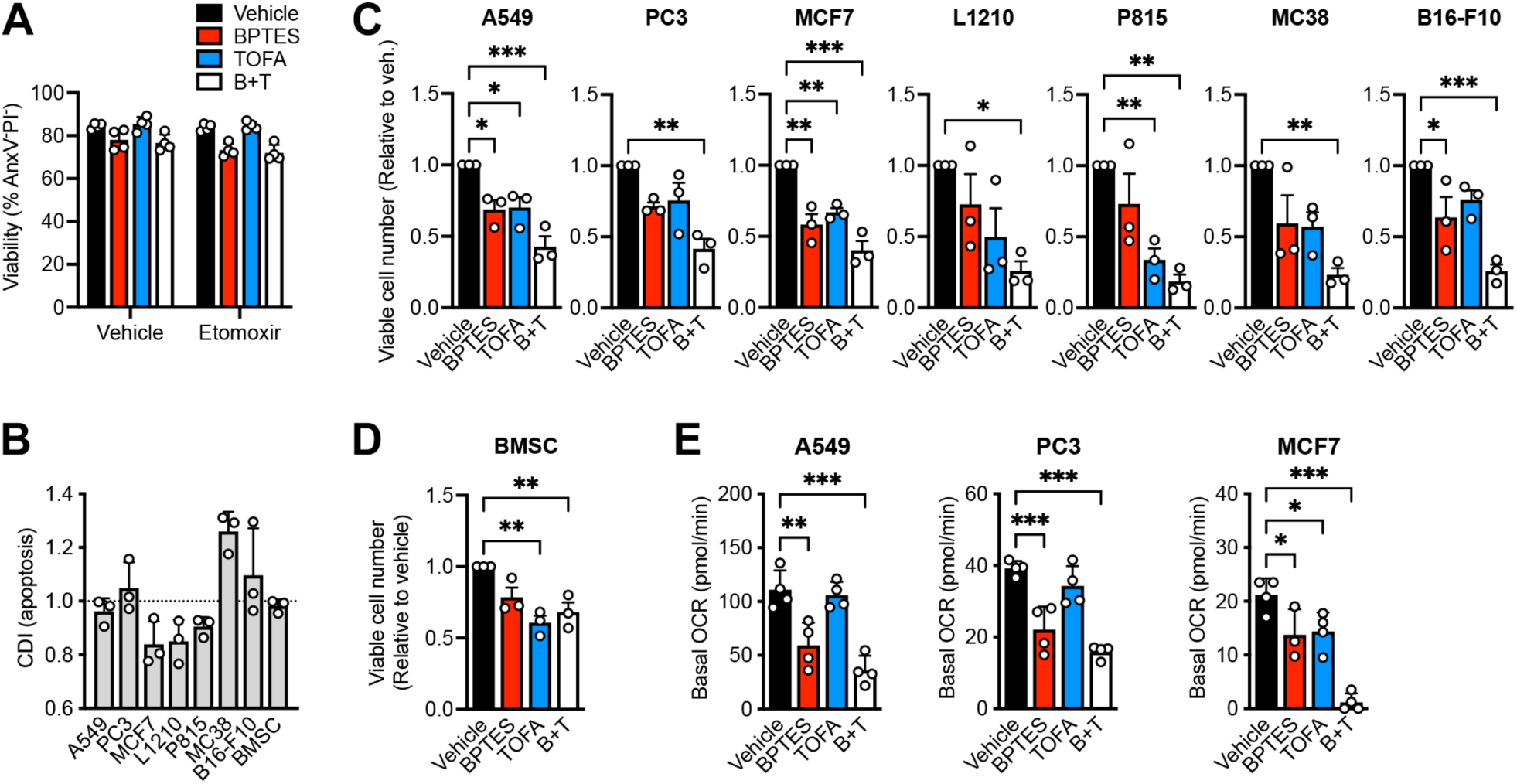
Cancer and Healthy Cells Show Different Responses to BPTES and TOFA (related to Figure 7). (**A**) Viability of cord blood-derived CD34^+^ cells treated for 72 hours with BPTES and/or TOFA in the presence or absence of etomoxir (40 µM). (**B**) Coefficient of Drug Interaction (CDI) for BPTES and TOFA calculated from the apoptosis data of different human and mouse cancer cell lines or healthy human bone marrow stromal cells (BMSC) shown in Figure 7I-J. (**C-D**) Viable cell number in cultures of different human and mouse cancer cell lines (C) or BMSC (D) treated for 72 hours with BPTES and/or TOFA. (**E**) Basal oxygen consumption rate (OCR) of three different human solid tumor cell lines treated for 24 hours with BPTES and/or TOFA, measured by Seahorse assay. *p < 0.05, **p < 0.01, ***p < 0.001.

