## Supplemental Table 1 for "Protein *S*-acylation dynamics provide metabolic plasticity to acute myeloid leukemia cells"

**Table S1. Genetic profile of AML patient samples and PDX lines.**

| Patient code | Genetic background | Treatment |
|---|---|---|
| AML160505B | FLT3 NM_004119 c.2028C>A p.N676K (31.6%)<br>IDH1 NM_005896 c.394C>T p.R132C (48.2%)<br>NPM1 NM_002520 c.859_860insTCTG p.W288fs*>9 (45.4%)<br>SRSF2 NM_001195427 c.284C>G p.P95R (52.9%) | Naïve |
| AML171026 | CBL NM_005188 c.1255T>A p.C419S (46.8%)<br>NPM1 NM_002520 c.859_860insTCTG p.W288fs*>9 (45.5%)<br>FLT3-ITD is detected | Naïve |
| AML171120 | DNMT3A NM_175629 c.2645G>A p.R882H (49.0%)<br>CEBPA NM_004364 c.940_941insAAG p.K313_V314insK (96.7%)<br>SETD2 NM_014159 c.914_915insA p.T305fs* (43.2%) | Naïve |
| AML180122 | PDS5B NM_015032 c.2163_2164insC p.R724fs*16 29.5%<br>RAD21 NM_006265 c.1090delG p.W363fs* 28.1% | Naïve |
| AML181002B | CEBPA NM_004364 c.877A>G p.N293D 26.8%<br>DNMT3A NM_175629 c.1627G>T p.G543C 43.6%<br>GATA2 NM_032638 c.959G>A p.G320D 36.7%<br>KIT NM_000222 c.2447A>T p.D816V 9.2%<br>TET2 NM_001127208 c.3312_3313insA p.F1104fs* 95.7%<br>U2AF1 NM_006758 c.101C>T p.S34F 48.7% | Naïve |
| AML181003B | GATA2 NM_032638 c.959G>A p.G320D 16.7%<br>RUNX1 NM_001754 c.496C>T p.R166* 43.5%<br>SF3B1 NM_012433 c.1998G>C p.K666N 41.0% | 22 cycles of azacitidine in treatment of prior MDS, MDS -> AML |
| AML181016B | PTPN11 NM_002834 c.178G>C p.G60R 7.6%<br>PTPN11 NM_002834 c.179G>T p.G60V 2.9%<br>PTPN11 NM_002834 c.205G>A p.E69K 3.3%<br>PTPN11 NM_002834 c.211T>C p.F71L 2.5%<br>PTPN11 NM_002834 c.213T>A p.F71L 8.9% | Naïve |
| AML181127D | ASXL1 NM_015338 c.1888_1910delCACCACTGCCATAGAGAGGCGGC p.E635fs*15 20.4%<br>PHF6 NM_001015877 c.823C>T p.R274* 38.0%<br>RUNX1 NM_001754 c.734_747delACACCACCCAGCCC p.H242fs*14 47.9%<br>SRSF2 NM_001195427 c.284C>G p.P95R 39.5% | Relapsed, post treatment |
| AML181214 | ASXL1 NM_015338 c.1888_1910delCACCACTGCCATAGAGAGGCGGC p.E635fs*15 31.2%<br>ASXL1 NM_015338 c.1902delA p.R634fs* 34.4%<br>EZH2 NM_004456 c.1979G>A p.G660E 45.3%<br>STAG2 NM_001042749 c.1798_1799insATACC p.Y600fs* 46.7%<br>TET2 NM_001127208 c.4521delG p.Q1507fs* 62.5% | No treatment in-house, may have been treated for prior MDS |
| DFAM-28971 | BCORL1 NM_021946 c.3490_3491insG p.R1164fs* 89.7%<br>DNMT3A NM_175629 c.2339A>C p.I780S 46.7%<br>NRAS NM_002524 c.182T>C p.Q61R 34.2%<br>NRAS NM_002524 c.35C>T p.G12D 2.5%<br>PTPN11 NM_002834 c.214G>A p.A72T 3.4%<br>TET2 NM_001127208 c.3176C>G p.S1059* 46.2%<br>U2AF1 NM_006758 c.101G>A p.S34F 35.3% | Naïve, prior hydroxyurea treatment |

| | | |
|---|---|---|
| MDAM-19692 | FLT3 ITD (ITD ratio is 0.467)<br>NPM1<br>DNMT3A<br>TP53 | Naïve |
| CBAM-68552 | MLL rearrangement at 11q23 confirmed by FISH | Relapsed, post chemotherapy |
