## Supplemental Table 4 for "Protein *S*-acylation dynamics provide metabolic plasticity to acute myeloid leukemia cells"

**Table S4. Mammalian enzymes using palmitoyl-CoA and/or stearoyl-CoA as substrate.**

| EC number | Enzyme activity | Human Gene(s) |
|---|---|---|
| 1.14.19.1 | stearoyl-CoA 9-desaturase | SCD, SCD5 |
| 1.2.1.84 | alcohol-forming fatty acyl-CoA reductase | FAR1, 2 |
| 1.3.3.6 | acyl-CoA oxidase | ACOX1, 3 |
| 1.3.8.8 | long-chain acyl-CoA dehydrogenase | ACADL |
| 2.3.1.15 | glycerol-3-phosphate 1-O-acyltransferase | GPAT1-4 |
| 2.3.1.20 | diacylglycerol O-acyltransferase | DGAT1-2 |
| 2.3.1.21 | carnitine O-palmitoyltransferase | CPT1A, CPT1B |
| 2.3.1.22 | 2-acylglycerol O-acyltransferase | MOGAT1-3 |
| 2.3.1.225 | protein S-acyltransferase | ZDHHC1-23 |
| 2.3.1.23 | 1-acylglycerophosphocholine O-acyltransferase | LPCAT1-4 |
| 2.3.1.24 | sphingosine N-acyltransferase | CERS1, 4, 5, 6 |
| 2.3.1.26 | sterol O-acyltransferase | SOAT1-2 |
| 2.3.1.291 | sphingoid base N-palmitoyltransferase | CERS5 |
| 2.3.1.299 | sphingoid base N-stearoyltransferase | CERS1 |
| 2.3.1.42 | glycerone-phosphate O-acyltransferase | GNPAT |
| 2.3.1.50 | serine C-palmitoyltransferase | SPTLC1-3 |
| 2.3.1.51 | 1-acylglycerol-3-phosphate O-acyltransferase | AGPAT1-5, LCLAT1 |
| 2.3.1.75 | long-chain-alcohol O-fatty-acyltransferase | AWAT1, AWAT2 |
| 2.3.1.76 | retinol O-fatty-acyltransferase | DGAT1-2, AWAT2 |
| 3.1.2.2 | palmitoyl-CoA hydrolase | ACOT1, 2, 4, 7, 8, 9, 11 |
| 7.6.2.4 | ABC-type fatty-acyl-CoA transporter | ABCD1-3 |
